# Fascetto Interacting Protein (FIP) Regulates Fascetto (PRC1) to Ensure Proper Cytokinesis and Ploidy

**DOI:** 10.1101/413997

**Authors:** Zachary T. Swider, Rachel K. Ng, Ramya Varadarajan, Carey J. Fagerstrom, Nasser M Rusan

**Author notes:** Corresponding Author Nasser M. Rusan. equal contribution (Zachary T. Swider, Rachel K. Ng).

## Abstract

Cell division is critical for development, organ growth, and tissue repair. The later stages of cell division include the formation of the microtubule (MT)-rich central spindle in anaphase, which is required to properly define the cell equator, guide the assembly of the acto-myosin contractile ring, and ultimately ensure complete separation and isolation of the two daughter cells via abscission. Much is known about the molecular machinery that forms the central spindle, including proteins needed to generate the antiparallel overlapping interzonal MTs. One critical protein that has garnered great attention is Protein Regulator of Cytokinesis 1 (PRC1), or Fascetto (Feo) in *Drosophila*, which forms a homodimer to crosslink interzonal MTs, ensuring proper central spindle formation and cytokinesis. Here, we report on a new direct protein interactor and regulator of Feo we named Fascetto Interacting Protein (FIP). Loss of FIP results in a significant reduction in Feo localization, rapid disassembly of interzonal MTs, and several cytokinesis defects. Simultaneous reduction in Feo and FIP results in tumor-like, DNA-filled masses in the brain. In aggregate our data show that FIP functions upstream of, and acts directly on, Feo to ensure fully accurate cell division.

## Introduction

Cell division, or mitosis, culminates in the separation of chromatin and cytoplasm into two daughter cells, in stages respectively known as anaphase and cytokinesis. These final stages of mitosis require exquisite coordination between three major cellular polymers – actin, microtubules (MTs), and septins. During anaphase, MT associated proteins (MAPs), molecular motors, and other effector proteins act on interdigitated, antiparallel MTs (referred to here as interzonal MTs) between opposite spindle poles to elongate the mitotic spindle and create the central spindle (D’Avino et al., 2015; Glotzer, 2016). The central spindle helps coordinate the position and assembly of the actin and myosin rich cytokinetic apparatus, or contractile ring, which is constructed in conjunction with the actin crosslinking protein Anillin and the septin cytoskeleton (Green et al., 2012; Liu et al., 2012; Maddox et al., 2007). The contractile ring then constricts around the central spindle which compacts into a midbody, or Flemming body. Finally, the midbody coordinates the abscission process that fully separates the two daughter cells (Mierzwa and Gerlich, 2014). Orchestrating anaphase, cytokinesis, and abscission is not only important for symmetric division, but is critical for asymmetric divisions where two non-identical daughter cells are produced, as is the case for stem cell divisions. Therefore, it is not surprising that defects in these final stages of mitosis have been linked to defects in cell ploidy, tissue development, and diseases such as cancer (Ganem et al., 2007; Lacroix and Maddox, 2012).

Given the importance of anaphase and cytokinesis, considerable attention has been given to these late mitotic processes, resulting in the identification of well conserved components that regulate the MT, actin, and septin networks. In search of novel regulators of cytokinesis, we investigated a set of proteins previously found to bind Septin 1 and Septin 2 in *Drosophila* using yeast 2 hybrid analysis (Y2H; Shih et al., 2002); they were termed Septin Interacting Proteins 1, 2 and 3 (Sip1, Sip2, Sip3). Sip2 was of particular interest as we found a striking localization to growing MT +ends and because it has not been investigated since its discovery. However, our preliminary work found no evidence of an interaction with Septins, or a direct role in regulating Septin function. Instead, we discovered that it binds another conserved cytokinesis molecule called Fascetto (Feo in *Drosophila*, PRC1 in mammals, Spd-1 in *C. elegans*) (Jiang et al., 1998; Verbrugghe and White, 2004; Verni et al., 2004), which binds and crosslinks MTs to form the central spindle (Mollinari et al., 2002; Subramanian et al., 2010). Thus, we have renamed Sip2 to Feo Interacting Protein (FIP) and focused our attention on how FIP functions as a Feo regulator.

A lot is known about Feo/PRC1 regulation throughout mitosis. Prior to anaphase, Feo/PRC1 is held inactive through inhibitory phosphorylation by Polo-like Kinase (Plk1; Hu et al., 2012; Neef et al., 2007), and/or Cdk1 (Wang et al., 2015; Zhu et al., 2006). Following anaphase onset, Feo/PRC1 forms a dimer (D’Avino et al., 2007; Kellogg et al., 2016; Subramanian et al., 2010), binds the Kinesin4/Kif4 (Klp3a in *Drosophila*), and localizes to the MT +ends (Bieling et al., 2010; D’Avino et al., 2007; Nguyen et al., 2018; Subramanian et al., 2013; Zhu and Jiang, 2005). Together, Feo/PRC1 and Kif4/Klp3a function to organize the central spindle in concert with Centralspindlin, a heterotetrameric complex comprised of a Kinesin-6 motor (MKLP in mammals, Pavarotti in *Drosophila*, ZEN-4 in *C. elegans*) and the RhoA GAP (RACGAP1 in mammals, Tumbleweed/RacGAP50C in *Drosophila*, CYK-4 in *C. elegans*) (Hirose et al., 2001; Jantsch-Plunger et al., 2000; Kurasawa et al., 2004; Mishima et al., 2002; Mishima and Lee, 2015; White and Glotzer, 2012). Feo/PRC1 is also involved in recruiting (directly or indirectly) additional proteins to the central spindle necessary for cytokinesis, such as Polo in *Drosophila* and Plk1 in mammals (D’Avino et al., 2007; Neef et al., 2007), the mammalian MT crosslinker Kinesin-5 (Subramanian et al., 2013), the *Drosophila* kinesin-14 and Sticky kinase (Bassi et al., 2013), mammalian CLASP1 (Liu et al., 2009), and mammalian CENP-E (Kurasawa et al., 2004). Thus, Feo/PRC1 is central to the overall organization of the complex protein networks required for daughter cell separation, and loss-of-function leads to mislocalization of MT associated proteins, thin/disorganized interzonal MTs, chromosome segregation errors such as chromosome bridge formation, incomplete furrow formation, cytokinesis failure, and ultimately aneuploidy and polyploidy (Li et al., 2018).

In our study, we show that FIP is a multifunctional adapter protein that exhibits several localizations *Drosophila*: MT +ends in interphase, the perichromosomal region in mitosis, interzonal MTs following anaphase onset, and at the midbody at the latest stages of mitosis. We show that loss of FIP results in defects similar to loss of Feo, and we provide both biochemical and genetic evidence to support a model in which FIP directly binds Feo to stabilize interzonal MTs needed for proper cytokinesis and genome stability.

## Results

### FIP is a microtubule and mitotic chromatin associated protein

To assess the intracellular localization of FIP, *Drosophila* S2 cells were transiently co-transfected with Ht::FIP (Ht = Halo-tag) and GFP::α-tubulin, fixed, and imaged by confocal microscopy. FIP localization changed dramatically throughout the cells cycle (Figure 1A). In interphase, FIP was enriched on MT +ends, suggesting that it may be involved in regulating MT dynamics. In prometaphase and metaphase, FIP was not detected on MTs, but clearly enriched around chromosomes, reminiscent of the perichromosomal sheath (Booth et al., 2016; Van Hooser et al., 2005). Following anaphase onset, FIP localized along interzonal MTs, which stimulate and stabilize the site of the cytokinetic-furrow. Finally, FIP decorated midbody MTs in late cytokinesis and through abscission, suggesting that FIP might play a role in the very final stages of cell division. To directly monitor FIP dynamics, we performed live 4D imaging of S2 cells expressing GFP::FIP. In agreement with fixed cell analysis, FIP localization rapidly changed from perichromosomal to the spindle midzone at anaphase onset, which then becomes compacted into the midbody MTs as cytokinesis progresses (Figure 1B; video 1).

**Figure 1.**
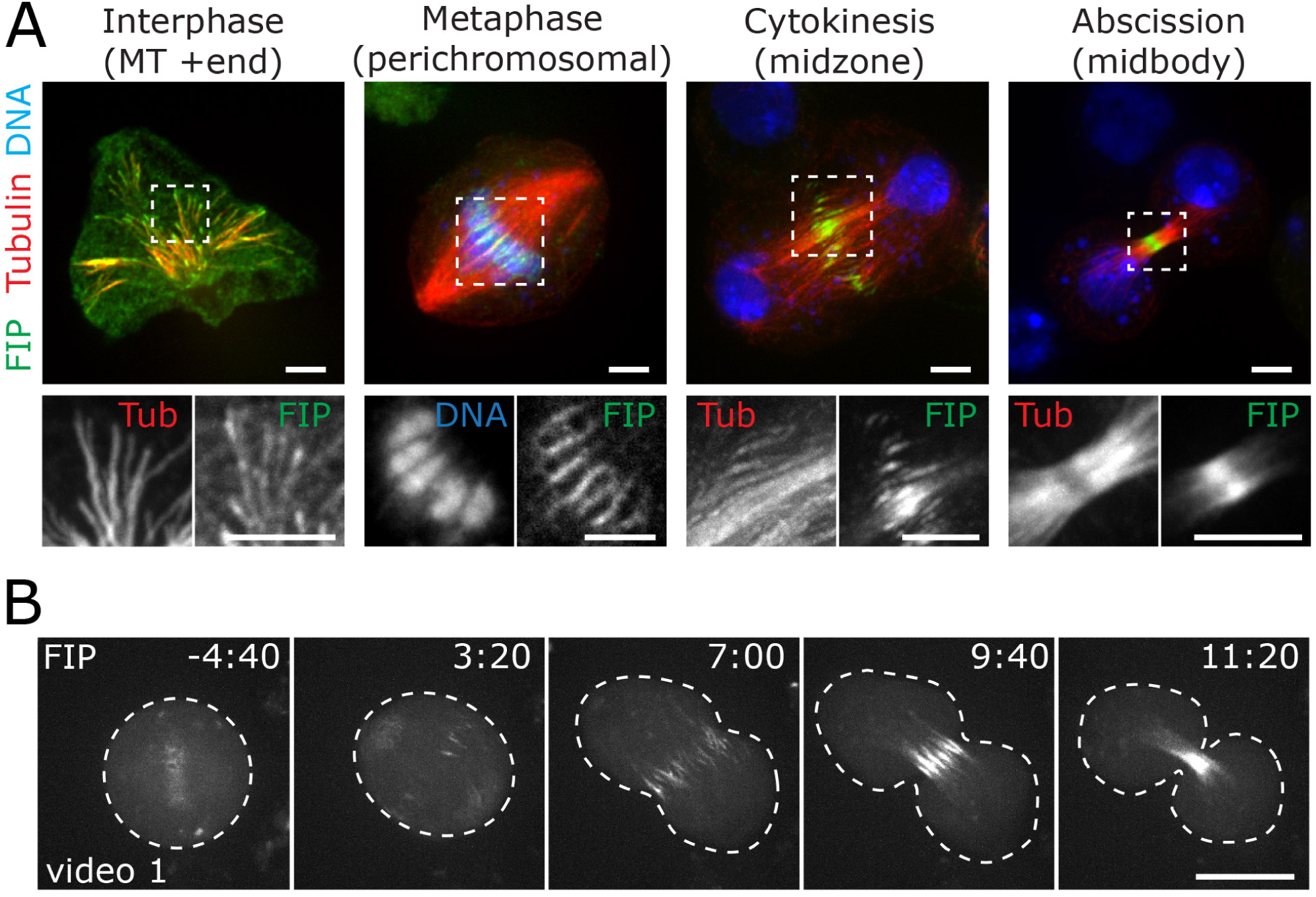
FIP shows cell-cycle dependent MT and chromatin dynamics. (A) Cell cycle stages of S2 cells coexpressing GFP::α-tubulin (Red) and Ht::FIP (Green; TMR ligand), MeOH fixed, and stained with DAPI (Blue). Regions outlined by dashed lines are enlarged in grayscale. Metaphase inset is a single image plane, which best represents FIP perichromosomal localization. All other images are maximum intensity projections. (B) Time series of a S2 cell expressing GFP::FIP (video 1) from metaphase to late telophase. Image stacks of six 0.8µm sections were collected every 20 sec and displayed as projections. Time = min:sec relative to anaphase onset. Scale bars: (A) = 5µm, (B) = 10µm.

The various subcellular localizations of FIP suggested that specific localization motifs or domains might be present; although no known protein domains are computationally predicted (Shih et al., 2002). To determine if such motifs were present, we truncated FIP into thirds, taking care not to disrupt predicted coiled-coil regions, and generated GFP fusions of each fragment (Figure 2A). The central region of FIP (FIP ^220-438^) is diffuse throughout the cytoplasm at all cell cycle stages (Figure 2B-D, center column), the N-terminus (FIP ^1-219^) conveys the perichomosomal localization in metaphase (Figure 2C, left column), and the C-terminus (FIP^439-657^) conveys localization to both interphase MT +ends and anaphase/telophase midzone MTs (Figure 2B, D, right column). Of note, FIP^439-657^ precociously localized to MT +ends in mitosis, indicating that upstream regions of FIP are required to prevent mitotic +end tracking (Figure 2C, right column).

**Figure 2.**
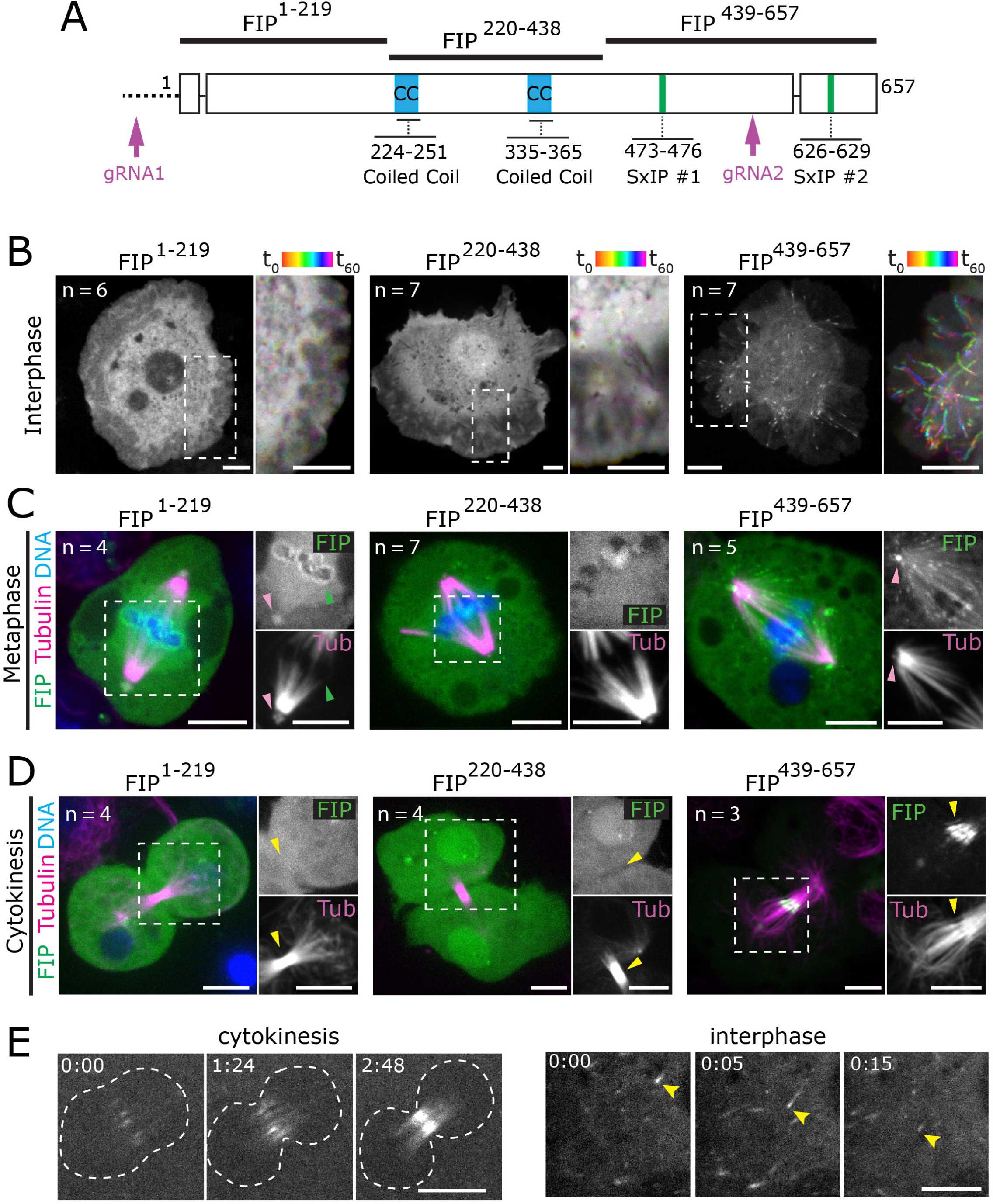
FIP subcellular localizations relies on specific protein regions. (A) FIP locus and protein-structure indicating the promotor region (dashed line), exons (boxes), introns (solid lines), predicted coiled coils (CC, blue), EB1 binding motifs (SxIP, green), the N-terminal, central, and C-terminal fragments used in this study (top, amino acid ranges indicated), and guide RNA sites used to generate the CRISPR mutant (magenta arrows; gRNA1 and gRNA2). (B) Interphase S2 cells expressing GFP-tagged FIP^1-219^, FIP^220-438^, or FIP^439-657^; stills from a 60 seconds time series (2 second intervals, single image plane; greyscale). Dashed lines represent the region of the cell used to create a color-coded ‘time projection’ (rainbow). Only FIP^439-657^ shows MT +end tracking (rainbow streaks). (C-D) Images of S2 cells in metaphase and cytokinesis expressing GFP-tagged FIP^1-219^, FIP^220-438^, and FIP^439-657^ (green) counterstained for DNA (Hoechst 33342, blue) and MTs (SiR-Tubulin, magenta). Regions with dashed lines are shown to the right in grayscale. FIP^1-219^ is perichromosomal (Green arrows) and centrosomal (Pink arrow). FIP^439-657^ localizes to centrosomes (Pink arrow) and midzone MTs during cytokinesis (Yellow arrow). All metaphase and cytokinesis images are maximum intensity projections through the region of interest. Ns on figure indicate number of cells imaged. (E) Wing disc cells expressing FIP::GFP show interzonal MT localization during cytokinesis and MT +end tracking in interphase. Scale bars: 5µm.

### FIP localizes to interphase MT +end through direct EB1-binding

To investigate FIP localization to MT +ends, we performed live 2-color imaging of S2 cells co-expressing fluorescently tagged FIP and the highly conserved MT <u>e</u>nd <u>b</u>inding <u>1</u> protein to mark MT +ends (EB1; Vaughan, 2005). In support of our fixed data, FIP co-localized with the characteristic EB1 MT +end-tracking comets in interphase cells (Figure S1A, video 2). FIP enrichment at MT +ends dropped from 1.46 ± 0.48-fold (over cytoplasm) in interphase to a nearly undetectable enrichment of 0.36 ± 0.24-fold in metaphase, while EB1 enrichment did not appear to change (Figure S1B-C, video 3). This regulated cell-cycle behavior is similar to the direct EB1-binding proteins STIM and CLASP2, which downregulate tip tracking behavior in response to mitotic phosphorylation (Kumar et al., 2012; Smyth et al., 2012). Thus, we hypothesized that FIP is also recruited to MT +ends by EB1. To test this hypothesis, we grossly overexpressed EB1 (EB1^OE^), a condition that drives exogenous EB1 along the MT lattice, and showed that FIP was also mis-localized to the MT lattice (Figure S1D). In contrast, overexpression of FIP alone never forced FIP on the MT lattice regardless of expression level (Figure S1E), indicating that EB1 is limiting for MT lattice recruitment.

Using yeast-2-hybrid (Y2H) analysis, we showed that both full length FIP (FIP^FL^) and the FIP C-terminus (FIP^439-659^) directly bind EB1 (Figure S1F). Additionally, FIP was co-immunoprecipitated from S2 cells overexpressing GFP::EB1 and Flag::FIP (Figure 6B). Sequence analysis of the FIP C-terminus revealed two consensus <u>S</u>er-<u>x</u>-Ile-<u>P</u>ro (SxIP) motifs (Figure 2A; Honnappa et al., 2009). To explore the function of these SxIP motifs, we mutated one (*fip*^*Δ1*^ or *fip*^Δ2^) or both (*fip*^*Δ1-2*^) SxIP motifs to SNNN (Jiang et al., 2012), and transfected these mutant constructs into S2 cells along with low levels of EB1. FIP^Δ1^ and FIP^Δ2^ exhibited a decreased enrichment at MT +ends, while *fip*^*Δ*^ was undetectable at +ends (Figure S2A-B, video 4). Furthermore, EB1^OE^ was unable to recruit *fip*^*Δ1-2*^ to the MT lattice (Figure S1G), confirming that the SxIP motifs mediate EB1-FIP interaction.

Given the MT +end localization, we tested if FIP plays a role in regulating interphase MT dynamics. Live imaging of interphase S2 cells depleted of FIP by dsRNA and expressing EB1-GFP revealed a slight, but statistically significant difference in MT growth rate (130 ± 65 nm/s compared to 150 ± 68 nm/s in controls; Figure S2C-D). Thus far, our results suggest that FIP is recruited to MT +ends by a direct interaction with EB1, but does not play a major role in MT growth. While we cannot rule out a role for MT rescue or catastrophe, it is unlikely given the absence of FIP on paused and shrinking MTs.

### FIP is required for efficient progression through mitosis

To investigate the role of FIP in a physiologically-relevant context, we turned to a detailed analysis of FIP in intact *Drosophila melanogaster* tissues. We generated transgenic flies expressing GFP::FIP driven by the ubiquitin promoter and imaged larval imaginal wing disc cells. Similar to in S2 cells, FIP localized to both midzone MTs during cytokinesis and MT +ends during interphase (Figure 2E). We then used CRISPR to generate *fip*^*7*^, a mutant allele that deletes 153 nucleotides of the promoter sequence, the transcription and translation start sites, and ∼75% (1452/1971) of the coding nucleotides (Figure 2A, gRNAs). Animals homozygous for *fip*^*7*^*/fip*^*7*^, or *fip*^*7*^*/Df(2L)Exel7029* (hereon both referred to as *fip-*) are viable and fertile; thus, FIP is a non-essential protein for viability of *D. melanogaster* reared in typical lab conditions. However, analysis of wing discs from identically aged third-instar *fip-* larvae were approximately 22 percent smaller with an average cross-sectional area of 0.18 ±.04 mm^2^ compared to 0.23 ±.03 mm^2^ in controls, indicating that FIP is required for proper tissue growth (Figure 3A).

**Figure 3.**
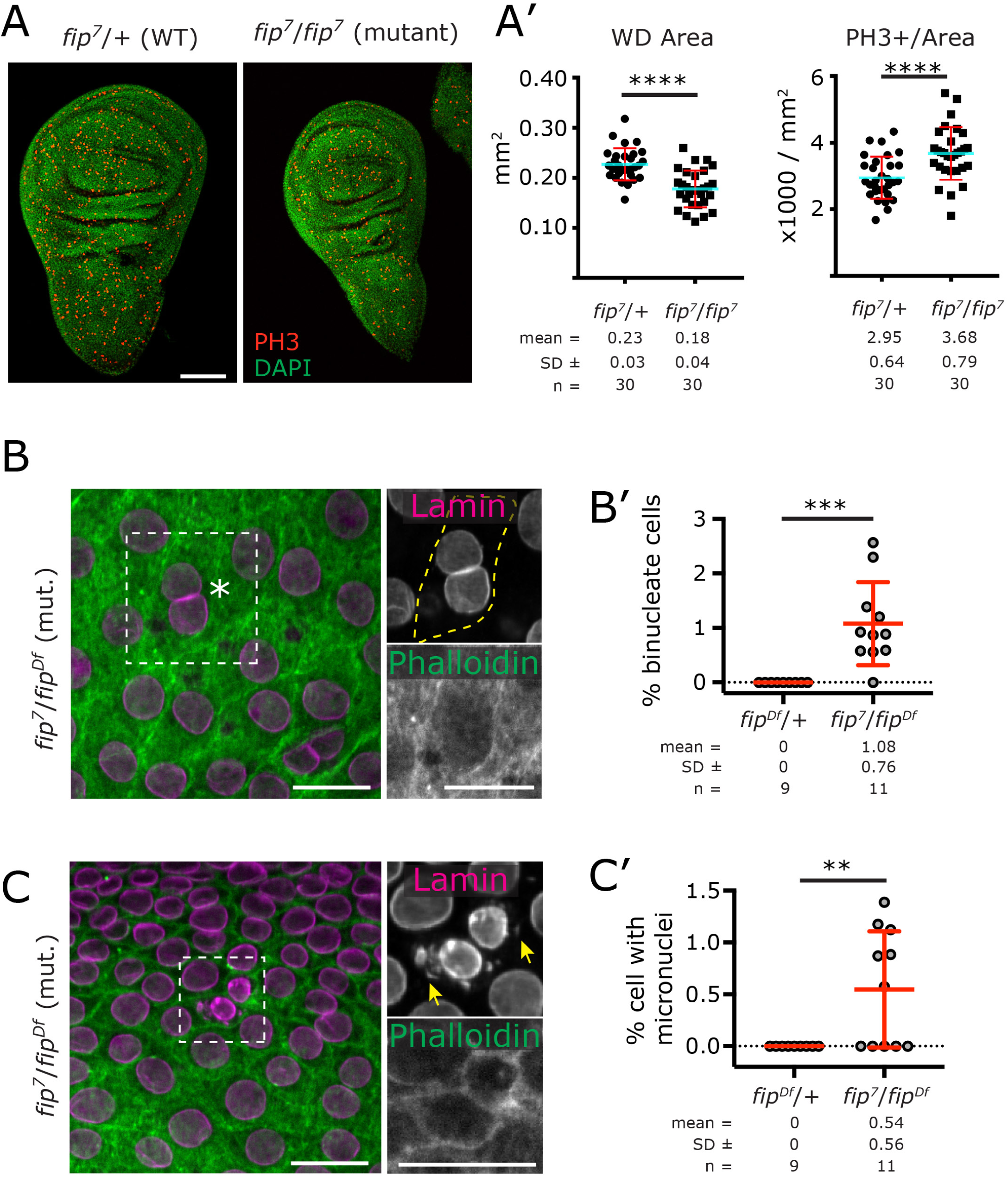
FIP is required for efficient cell division. (A) Wing disc from wt and *fip-* 3^rd^ instar larvae dissected 96 hours after egg laying stained for DAPI (green) and PH3 (red). (A’) Wing disc area and number of PH3+ cells, normalized to area as a proxy for cell number. Data from 5 independent experiments, **** = P < 0.0001. (B) Binucleate cell in a fixed *fip-* wing disc (dashed box, asterisk) stained with phalloidin (green) and anti-lamin (magenta). The region within the dashed box is shown in grayscale on the right. (B’) Percentage of cells that were binucleate; each point represents a single wing disc, *** = P < 0.001. (C) Micronuclei in a fixed *fip-* wing disc (dashed box) stained with phalloidin (green) and anti-lamin (magenta). The region within the dashed box is shown in grayscale on the right. (C’) Percentage of cells with micronuclei; each point represents a single wing disc, ** = P < 0.01. Scale bars: (A) = 100µm, (B, C) = 5µm

Given the localization of FIP in mitosis, we predicted that the growth defect was due to a mitotic defect. Indeed, analysis of fixed *fip-* wing discs showed an elevated mitotic index (3.68×10^3^ ±.79×10^3^PH3 positive cells/mm^2^compared to 2.95×10^3^ ±.64×10^3^in controls; Figure 3A, A’), binucleate cells (1.08 ± 0.76% of cells; Figure 3B, B’), and rare incidences of micronuclei (0.54 ± 0.56% of cells; Figure C, C’). These data revealed a history of cytokinesis failure and possibly chromosome fragmentation or mis-segregation (Fenech et al., 2011), which we attempted to capture using two color live imaging of mitosis in *fip-* wing discs using GFP::Jupiter (marking MTs) and H2AV::mRFP (marking chromosomes). Although we did not capture complete mitotic failure, our live imaging uncovered a slight delay in mitotic progression (nuclear envelope breakdown (NEBD) to anaphase onset of 533 ± 116 seconds in mutants compared to 493 ± 67 seconds in controls; Figure 4A, A’) and a defect in chromosome segregation wherein *fip-* cells ceased chromosome movement sooner than controls and segregated a shorter distance (Figure 4B, B’). Parallel experiments using dsRNA knockdown of FIP in S2 cells revealed multinucleate cells (6.2 ± 2.5%, vs 3.7 ± 2.0% in controls) and a decrease in anaphase cell index (0.1 ± 0.0%, vs 0.7 ± 0.8% in controls) (Figure S2E). Taken together, these results suggest FIP is required for fully accurate chromosome segregation, cytokinesis, and overall mitotic progression.

**Figure 4.**
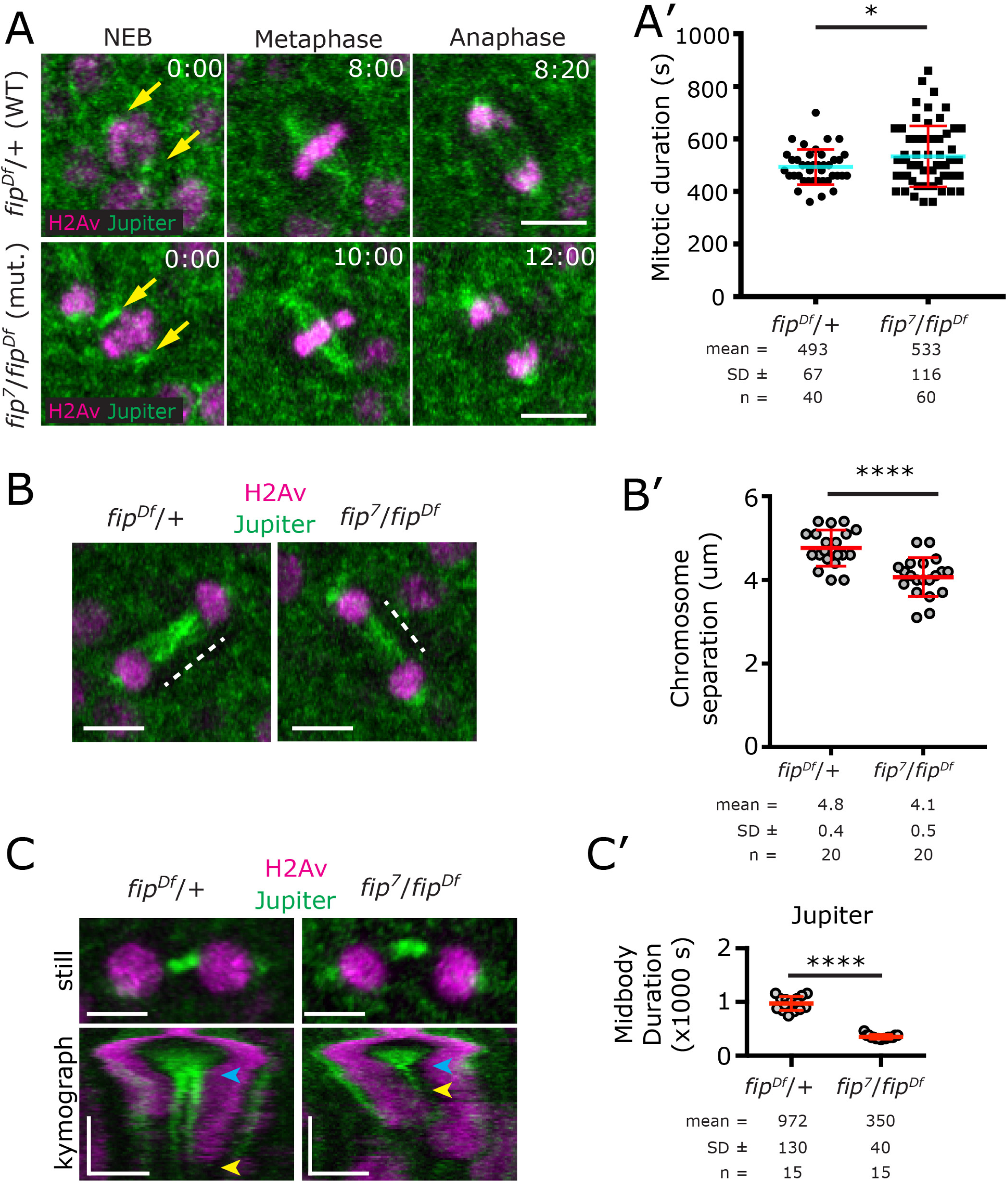
FIP is required for interzonal MT stability. (A) 4D time series of control and *fip-* wing disc expressing GFP::Jupiter (green) and H2Av::mRFP (magenta) progressing through mitosis. (A’) Quantification of mitotic duration (NEBD to anaphase onset) showing a delay in mitotic progression in *fip-* cells, * = P < 0.05. (B) Wing disc cells in telophase showing maximum nuclear separation. Dashed lines represent chromosome separation measurements. (B’) Quantification of maximum chromosome separation (displayed as % of metaphase spindle length to control for cell size); each point represents a single cell, **** = *P* < 0.0001. (C) Stills and kymographs from telophase *fip-* and control wing discs. Kymographs begin at anaphase (separation of chromosomes, magenta streaks). Wild-type midbody formation in controls is seen as a pair of streaks (green), while the rapid dissolution of MTs (GFP::Jupiter) is seen in the *fip-* kymograph. Blue arrows indicate the frame of the kymograph used to create the still image above, yellow arrows indicate where the GFP::Jupiter interzonal signal ends. (C’) Interzonal MT duration measured using GFP::Jupiter signal (anaphase onset to the last frame where an intercellular MT bridge was clearly seen); each point represents a single cell, **** = *P* < 0.0001. Horizontal scale bars in (A) (B) (C) = 3µm, vertical scale bars in (C) = 480 seconds.

In searching for the mechanism by which loss of FIP could cause these defects and considering previous work that showed FIP binds Septin 1 and Septin 2 (Shih et al., 2002), we hypothesized that FIP regulates septins to ensure proper cytokinesis. Indeed, loss of septin family members has been shown to result in cytokinesis failure (Kinoshita et al., 1997; Nagata et al., 2003), and chromosome missegregation (Spiliotis et al., 2005). We were unable, however, to confirm direct FIP-Septin interaction by Y2H (Figure S3A), and our two color live imaging of Sep2::GFP and H2AV::mRFP in dividing *fip-* wing disc cells revealed that the timing of Septin 2 recruitment to the cleavage furrow and midbody appeared normal (data not shown). Most interestingly, we found that Septin 2 was less persistent at the midbody (disassembling in 2002 ± 330 seconds following anaphase onset compared to 2784 ± 623 seconds in controls; Figure S3B, B’), and high-resolution imaging in fixed *fip-* cells revealed that Septin 2 in the midbody lacked the cylindrical organization characteristic of control midbodies (Figure S3C). Given that FIP does not localize to the cell cortex and that we were unable to detect direct FIP-Septin interaction, we favor the hypothesis that the defects in Septin organization and dynamics were secondary to an upstream defect.

### FIP stabilizes interzonal MTs during cytokinesis

Based on FIP’s localization to interzonal MTs, we hypothesized that the atypical organization of Septin 2 in *fip-* midbodies was due to defects in MT organization. Our live imaging of GFP::Jupiter (marking MTs) and H2AV::mRFP in *fip-* wing discs revealed no significant defect in interzonal MT density in *fip-* wing discs in the first two minutes following anaphase onset (measured by GFP::Jupiter enrichment, not shown). Following *fip-* wing disc cells further into telophase revealed a prominent defect where interzonal MTs focused into a single point, in contrast to wild-type cells that show a stereotypical organization of two discrete bundles of MTs on either side of the midbody (Figure 4C, parallel vertical streaks in the kymograph). It appears as if the antiparallel, overlapping interzonal MTs region is significantly diminished, possibly completely lost. Even more striking, however, was that *fip-* cells lost all detectable midbody MTs three times faster than controls (Figure 4C, yellow arrows; 4C’, 350 ± 40 seconds after anaphase onset compared to 952 ± 130 seconds in controls).

Having documented defects in both the organization and stability of MTs and failures in cytokinesis when FIP is absent, we hypothesized that loss of FIP would consequently lead to defects in the actin contractile ring structure. To test this, we indirectly followed cortical actin by imaging GFP::Moesin and found that 100% of *fip-* cells completed the constriction of the cytokinetic ring down to the midbody (*n* = 120 cells from 3 wing discs) with no significant difference in the timing (154 ± 26 seconds from furrow initiation to completion compared to 153 ± 25 seconds in controls; Figure S4A, B), no significant difference in actin enrichment in the nascent furrow (2.62 ± 0.5-fold enrichment compared to 2.44 ± 0.5-fold in controls; Figure S4C), and no significant difference in the duration of GFP::Moesin localization to the midbody (882 ± 127 seconds compared to 986 ± 229 in controls; Figure S4D). Together, these results suggest that FIP is dispensable for early organization of interzonal MTs and actin ring formation but plays an important role in stabilizing the late interzonal MTs required for the very final stages of cell division.

### FIP is required for proper cell ploidy and brain development

Previous studies have shown that mutations in mitotic genes can result in variant phenotypes in the wing disc and larval brain – in some cases, mutant brains exhibit a higher rate of polyploidy (Poulton et al., 2014; Poulton et al., 2017). Indeed, analysis of *fip-* larval brains revealed striking chromatin masses (Figure 5A, B; 4.5± 4.5 polyploid cells per central nervous system (CNS) compared to 0/CNS in controls). Expression of GFP::FIP in the *fip-* background nearly fully recued the polyploid phenotype (1.1± 1.9 polyploid cells per CNS; Figure 5B). High resolution imaging of these chromatin DNA masses showed that they typically reside within single cells, adopting a lobular morphology often surrounded by numerous micronuclei (Figure 5C, red arrows).

**Figure 5.**
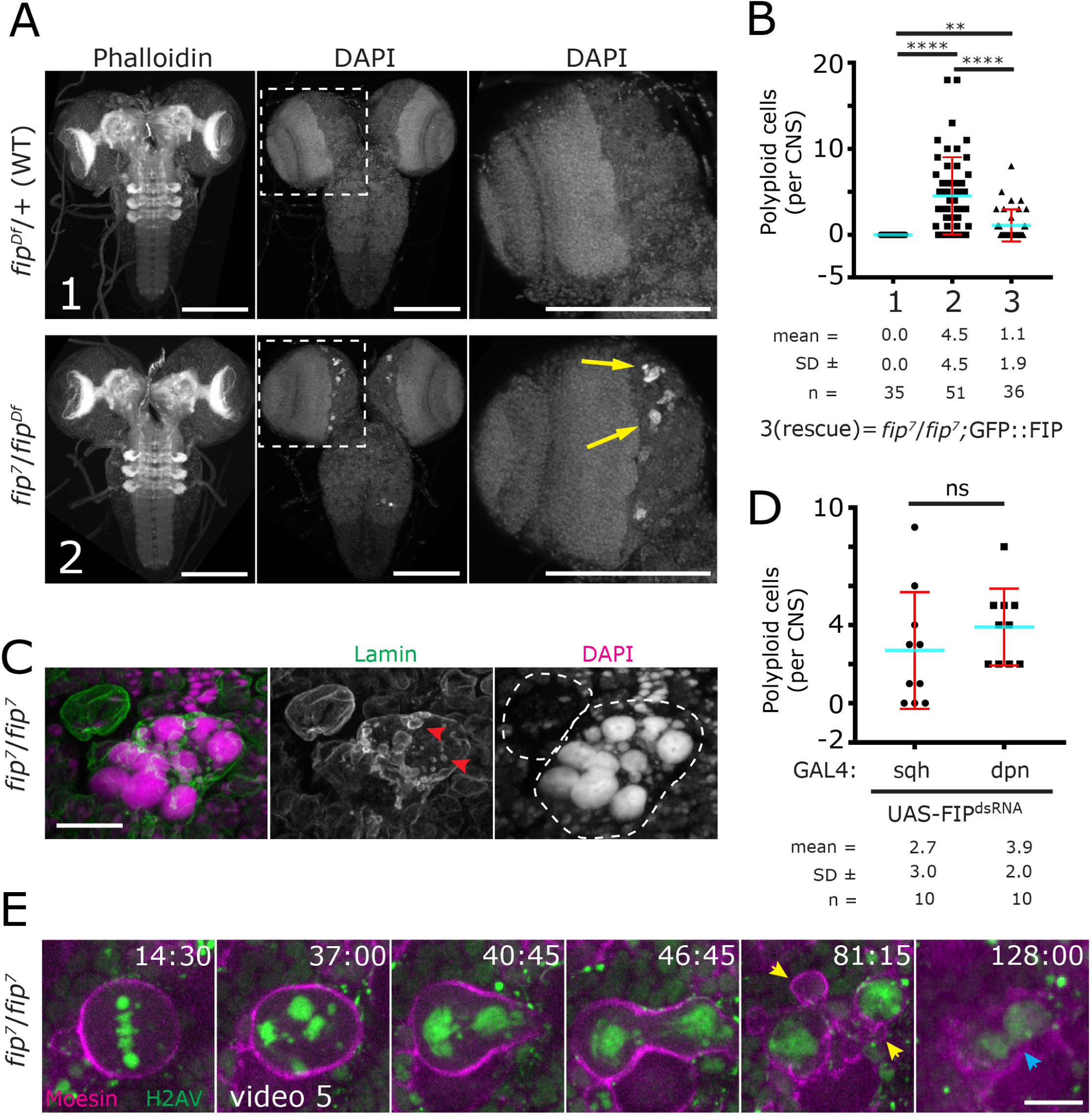
Loss of FIP results in failed divisions and polyploidy in brains. (A) Third instar larval brains stained for F-actin (phalloidin) and DNA (DAPI). Dashed boxed region is enlarged to the right, highlighting polyploid neuroblasts (NBs) in the *fip-* mutant(yellow arrows). (B,D) Quantification of the number of polyploid cells in control, mutant, rescue, and RNAi knockdown conditions; each point represents the number of polyploid cells per centralnervous system (CNS = central brain + ventral nerve cord (VNC)) sqh-Gal4 is ubiquitous RNAi knockdown, Dpn-Gal4 is NB specific. The numbers on the x-axis (1,2,3) refer to the genotype indicated in A or below the graph. (C) High resolution image of a polyploid cell adjacent to a healthy NB (top left) stained with anti-lamin (green) and DAPI (magenta) showing many large DNA structures in addition to micronuclei (red arrows). Dashed lines indicate cell boundaries from phalloidin channel (not shown). (E) Live image of a *fip-* NB expressing GFP::Moesin (magenta) and H2AV::mRFP (green), see video 5. NB initiates asymmetric division (40:45), attempts to complete cytokinesis (46:45), begins massive membrane blebbing at the site of furrow formation (81:15, yellow arrows), and eventually fails cytokinesis/abscission, regressing into a single cell (128:00, blue arrow). Time is in min:sec relative to NEB. Scale bars: (A) = 150µm, (C) and (E) = 10µm.

The distinctive localization of the polyploid cells in the CNS suggested that these cells were derived from central brain neuroblasts (NBs). To explore this possibility, a UAS-FIP RNAi transgene was driven by either *Sqh*-GAL4 (a strong and ubiquitous driver) or *Dpn*-GAL4 (a relatively moderate driver expressed only in NBs). Both gene knockdowns produced identical polyploid cells in the central brain (2.7 ± 3 /brain in *Sqh*-GAL4 and 3.9± 2 /brain in *Dpn*-GAL4; Figure 5D). To better understand how the polyploidy arises, we performed live two-color imaging of GFP::Moesin and H2AV::mRFP in *fip-* brains. We were unsuccessful in capturing the initial event of polyploidization; however, we did capture an early polyploid NB in mitosis, which remained in metaphase for ∼33 minutes (compared to the published average 8 minutes, Rusan et al., 2008) with several misaligned chromosomes (video 5). The NB eventually entered anaphase with chromatin bridges remaining in the division plane, ultimately failed abscission, and forced the cell into a greater state of polyploidy. We suspect that once a *fip-* NB undergoes a mitotic failure, it progressively becomes worse with each round of attempted division.

### FIP binds the MT lattice via the PRC1 ortholog Fascetto (Feo)

To explore the mechanism by which FIP regulates cytokinesis, we sought to identify protein partners of FIP that might function at interzonal MTs. Although we showed that FIP localization did not appear exclusive to MT +ends in the later stages of mitosis, we nevertheless tested if FIP localization to interzonal MTs was mediated by EB1. Clearly, however, live imaging of the EB1-binding mutant of FIP (*fip*^*Δ1-2*^) revealed normal interzonal MT localization in anaphase and telophase (Figure S4E, video 6); thus, interzonal MT recruitment is independent of EB1. We next investigated if FIP directly interacted with the MT binding protein Abnormal Spindle (Asp), which displays a similar anaphase localization to FIP (Asp; Riparbelli et al., 2002; Wakefield et al., 2001) and is known to bind MTs (Ito and Goshima, 2015; Mollinari et al., 2002; Saunders et al., 1997; Schoborg et al., 2015). However, we were unable to confirm a previously reported FIP-Asp interaction (Asp; Giot et al., 2003) using our Y2H system (Figure S5). Furthermore, GFP::FIP localization in a *asp-* genetic background appeared unperturbed (not shown). Lastly, we hypothesized that FIP is recruited to the central spindle by Fascetto (Verni et al., 2004), the *Drosophila* PRC1 ortholog shown to play an interzonal MT stabilization role during anaphase (Wang et al., 2015). We first used Y2H to reveal that full length FIP and Feo directly interact (Figure S5). Y2H analysis using FIP and Feo truncations narrowed the interaction down to FIP^220-647^(containing the sequence necessary and sufficient for MT localization in mitosis) and Feo^1-346^(containing the dimerization and rod domains) (Figure 6A). We also confirmed the interaction by co-immunoprecipitation from S2 cells overexpressing GFP::Feo and Flag::FIP (Figure 6B, S6A). Importantly, a Feo-FIP IP was only successful when the culture was enriched for mitotic cells using an overnight colchicine treatment, indicating the interaction is likely MT-independent and restricted to mitosis. This is consistent with the localization of Feo and FIP to separate cellular compartments in interphase – Feo is nuclear and FIP is cytoplasmic. As a final confirmation of FIP-Feo interaction *in vivo*, we performed live imaging of S2 cells co-expressing mNeonGreen::Feo and TagRFP::FIP. In low expressing interphase cells, FIP tracked MT +ends and Feo was seen along a subset of bundled MTs (Figure S6B, video7), while in high expressing cells we found that Feo ubiquitously coated and bundled MTs, as expected given Feo’s known role to crosslink MTs in mitosis (Figure S6B; Mollinari et al., 2002). In addition, these bundled MTs recruited FIP, which in interphase is found only at the MT +ends regardless of expression level (Figure S1E), indicating that Feo can mediate a FIP-MT interaction (Figure S6B). At low expression levels, and in the physiologically relevant context of mitosis, Feo and FIP localization were indistinguishable (Figure 6C, video 8). In *Drosophila* larval wing dics, Feo localization was also identical to FIP, showing translocation from the central spindle in anaphase to the midbody in telophase (Figure S7). Together, these results indicate that Feo and FIP likely form a complex and co-target interzonal MTs at anaphase onset.

**Figure 6.**
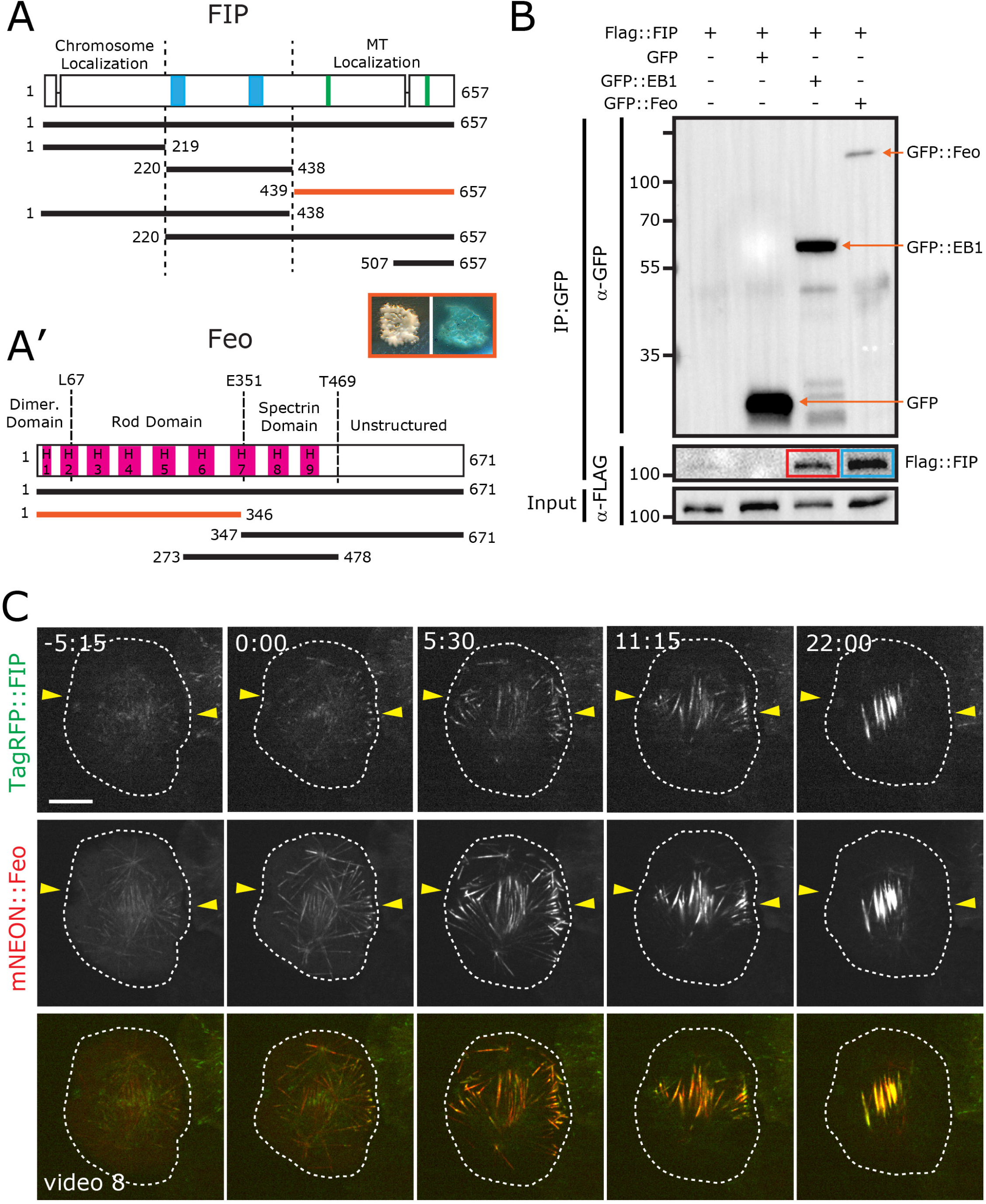
FIP directly binds Feo. (A) The indicated FIP protein truncations (horizontal lines) were tested for direct binding to Feo protein truncations (A’) via Y2H. The MT-localization region (439-657) of FIP binds to the N-terminal region (1-346) of Feo, which consists of the dimerization (1-67) and rod (68-351) domains. The blue yeast colony indicates a detectable interaction (growth is on QDOXA plates) between the two minimum fragments (orange lines). Complete interaction data is provided in Figure S5. (B) Western blot showing FIP co-immunoprecipitated with both GFP::EB1 (from asynchronous cells, red box) and GFP::Feo (from mitotically enriched cells, blue box). (C) S2 cells co-expressing mNeonGreen::Feo (red) and TagRFP::FIP (green) show identical localization of Feo and FIP beginning at anaphase onset (0:00) through telophase (22:00). Cell fails cytokinesis because it is plated on Con-A (video 8). Yellow arrows indicate the approximate position of the cell equator. Scale bar: (C) = 5µm.

### Feo requires FIP for proper localization

To test how FIP might regulate Feo localization, we turned to wing discs dissected from third instar larvae because of the abundance of mitotic cells and the ease of imaging. Wing discs from wild-type and *fip-* animals expressing Feo::GFP were fixed and stained for the contractile ring component Peanut (Pnut, Septin 3). Wild-type wing discs contain a high percentage of mitotic cells, with anaphase/telophase cells showing a clear contractile ring with Feo decorating interzonal MTs which compact down to a barrel-like midbody at the end of mitosis. In contrast, fixed *fip-* wing discs show nearly undetectable Feo on midzone MTs during anaphase and telophase (Figure 7A, B). At later stages of telophase and midbody formation, Feo levels are extremely low, and in 19% of midbodies (shown by Pnut) no Feo::GFP is detected. Interestingly, the low levels of Feo in *fip-* cells present as small dots (Figure 7B), mirroring the observation that MTs focus down to a small dot in *fip-* cells (Figure 4C). We also imaged these same wing discs live, which more clearly show the difference in mitotic activity and Feo levels between wildtype and *fip-* tissue (video 9, 10). However, we note that in contrast to our fixed analysis, live imaging of *fip-* wing discs show Feo::GFP on some interzonal MTs in anaphase and telophase (video 10). We suspect that Feo localization in *fip-* wing discs does not survive our fixation protocol possibly because the MTs are less stable and are hyper sensitive to the cold PBS fixation protocol required to detect Pnut (methods). However, even when we repeated the fixation with warm PBS, Feo::GFP was still undetectable on interzonal MTs (not shown). Taken together, these data indicate that FIP is required for normal levels of Feo localization and stabilization of interzonal MTs.

**Figure 7.**
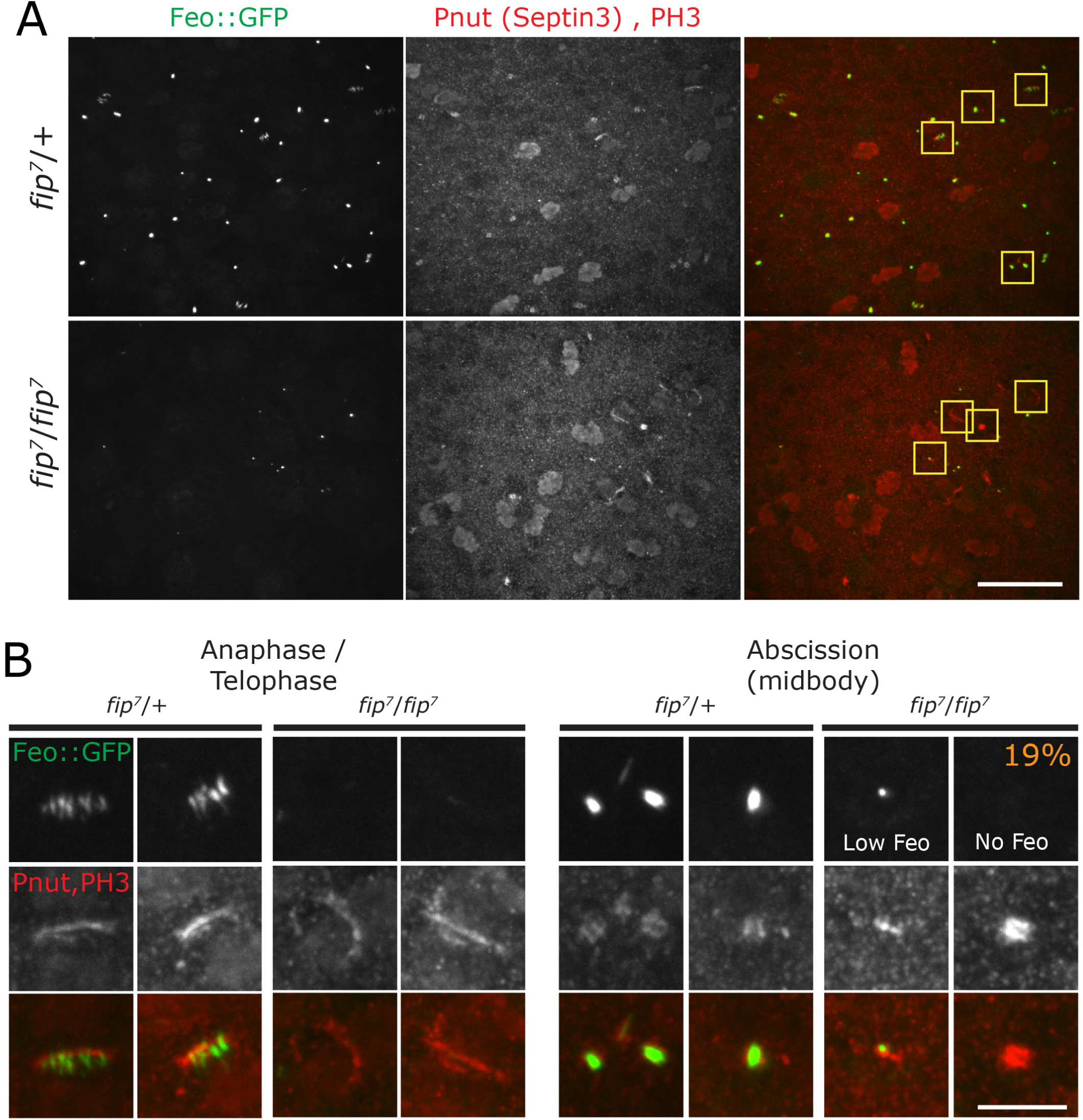
FIP is required for FEO localization and proper midbody architecture. (A and B) Peripodial cells in the pouch of control and *fip-* wing discs expressing Feo::GFP (Green) and stained for Pnut (Septin 3, Red) and PH3 (Red). (B) Enlarged images of peripodial cells show Feo::GFP localization at the anaphase spindle in control but not in *fip-* mutants. Feo prominently localizes to the midbody in controls while *fip-* mutant have low levels of Feo that appear focused down to a single dot; in 19% of cells, Feo is completely absent. Scale bars: (A) = 20um, (B) = 5um

### FIP and Feo function in the same pathway to ensure proper cell division

Having established a direct protein interaction, identical intracellular localization, and the localization dependence of Feo on FIP, we investigated a genetic interaction between FIP and Feo. For this analysis, we turned to the most prominent phenotype of polyploidy in the central brain (Figure 5). *fip-* (*fip*^*7*^*/fip*^*7*^ or *fip*^*7*^*/fip*^*Df*^) and *feo*^*RNAi*^ (tubulin-Gal4 driving *feo*^*RNAi(GL)*^ shown, or *feo*^*RNAi(HM)*^ not shown) brain lobes contain 1.9±2.1 and 0.6±.9 polyploid cells per lobe respectively (Figure 8A, A’), all within the central brain. *feo*^*RNAi*^ was used in this experiment because animals null for *feo* die at early larval stages. Importantly, double loss-of-function analysis (*fip-*; *feo*^*RNAi*^) showed a great enhancement of the phenotype with 8.4±3.6 polyploid cells per brain lobe (Figure 8A, A’). Animals co-depleted of FIP and Feo also greatly expand the effected brain tissue beyond the central brain and into the optic lobes (Figure 8A, blue arrows). Fly survival is also affected as loss of both FIP and Feo result in pupal lethality, whereas *fip-* or *feo*^*RNAi*^ alone give rise to viable adults. Finally, overexpression of Feo::GFP in the *fip-* mutant background nearly fully rescued the phenotype with an average of 0.3±.8 polyploid cells per lobe (Figure 8A, A’). Similar results to the brain lobes were observed when analyzing cells in the ventral nerve cord (VNC; Figure 8B, B’). Together these results argue that FIP and Feo function together to ensure proper cytokinesis.

**Figure 8.**
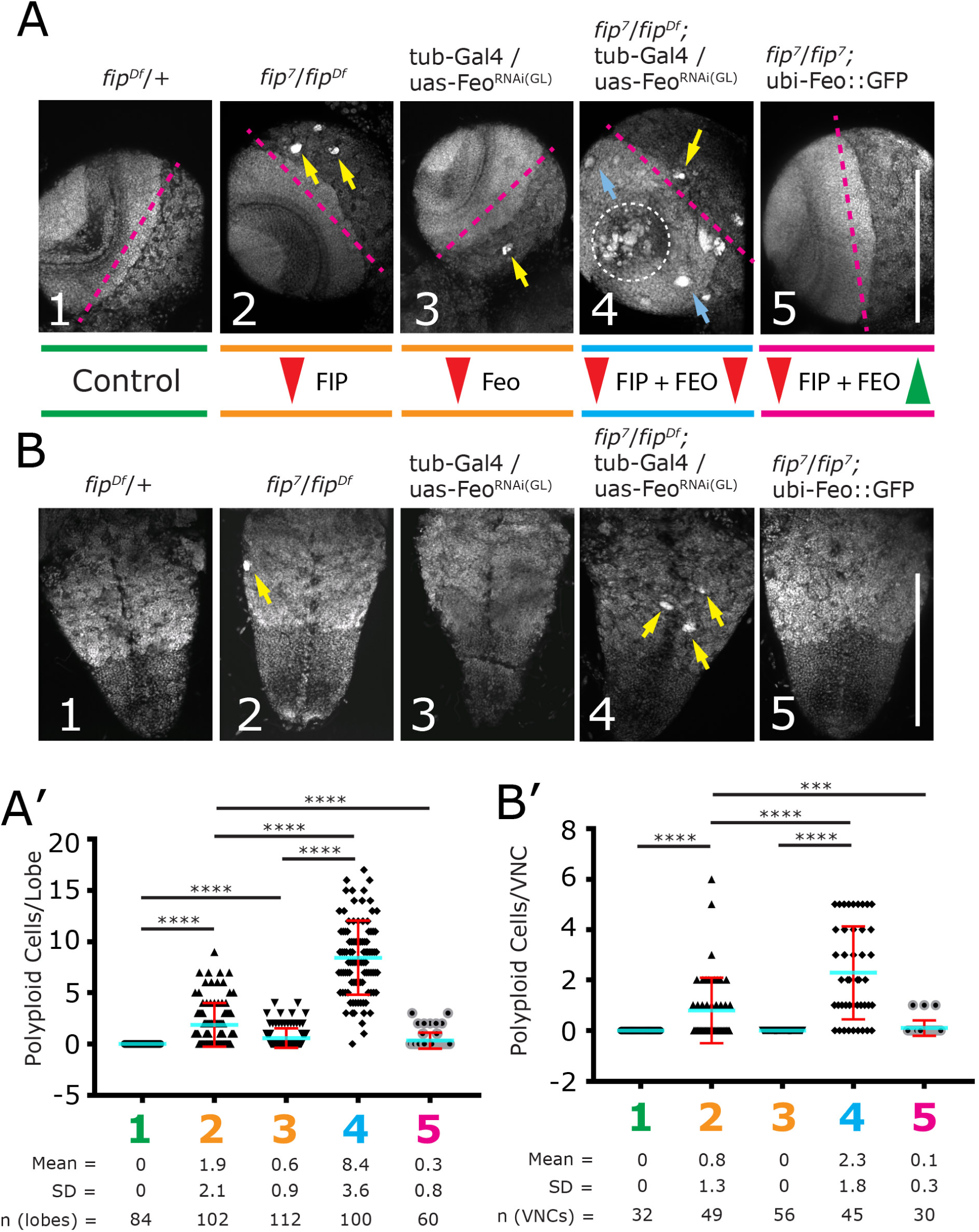
FIP and Feo function in the same pathway to ensure proper brain cell ploidy. (A) Third instar larval brains stained for DNA (DAPI) in control (Green, 1), single loss-of-function (Orange, 2 and 3), and double loss-of-function (Blue, 3). Overexpressing Feo in the *fip-* background (Pink, 5) rescues the polyploidy phenotype. Red arrows indicate loss of function; Green arrows indicate overexpression. Pink dashed lines roughly delineate the central brain (right) from the optic lobe (left). Polyploid cells in the central brain (yellow arrows) are present in single loss-of-function while polyploid cells are present in both the central brain (yellow arrows) and optic lobes (blue arrows) in double loss-of-function. (A’) Quantification of polyploid cells per brain lobe. (B) Ventral nerve chord (VNC) stained for DNA (DAPI), genotypes as indicated above. (B’) Quantification of polyploid cells in the VNC. Scale bar: 150um.

Interestingly, a more careful analysis using higher magnification of the polypoid cells in either *fip-* or *feo*^*RNAi*^ revealed a variety of nuclear morphologies, such as clusters of small DNA fragments and larger DNA masses contained within a single cell (Figure 9A, representative samples). In contrast, double loss-of-function (*fip-*; *feo*^*RNAi*^) resulted in 64% of brains containing much larger DNA structures (Figure 9B), in addition to the previously mentioned increase in overall number of polyploid cells (Figure 8A, A’). These structures can be grouped into two categories – ‘Single Mass’ DNA, which are homogenous DNA structures contained within a single huge cell (Figure 9C), and ‘Clustered’ DNA fragments (Figure 9D). To convey the difference in size, we measured the cross-sectional area of the DNA regions in wild-type and in single and double loss of function (Figure 9E). The average cross-sectional area of a wild-type NB nucleus was 100 ± 5 um^2^, while the average size of a *fip-* polyploid nucleus was 570 ± 130um^2^, and a *feo*^*RNAi*^ polyploidy nucleus was 550 ± 140 um^2^. The average size of the large ‘single masses’ in the double loss-of-function was nearly 50-times larger (4742 + 1821 um^2^) than wild-type, while the ‘clustered’ nuclei were over 100-times larger (10147 + 2815 um^2^). These larger masses could be a result of cytokinesis failures earlier in development, or a group of many cells that failed cell division and then merged into a DNA aggregate. Together, these results support a model where FIP is positioned upstream of Feo in a genetic pathway that regulates cytokinesis.

**Figure 9.**
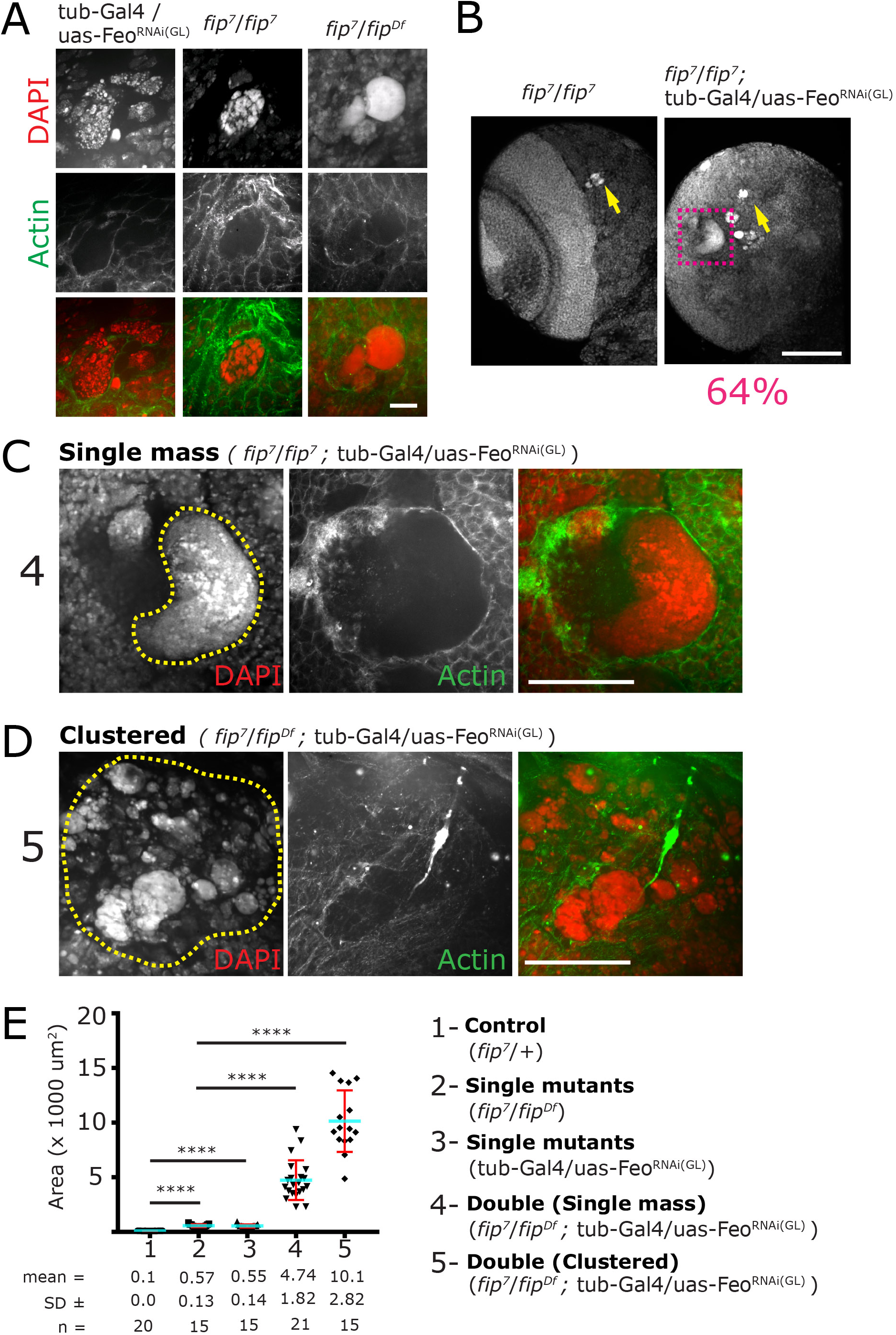
Enhanced polyploid and large DNA masses form in *fip-* and Feo^RNAi^ co-depleted animals. (A) Representative images of polyploid cells in *fip-* and Feo^RNAi^ brains. Polyploid cells have varied DNA morphologies including fragmented nuclei and dense DNA aggregates. (B) The single *fip-* mutants show relatively small polyploid cells, while the double depletion of *fip-* and Feo form massive DNA aggregates that span all brain regions. (C and D) The large DNA aggregates were generally classified into ‘single’ and ‘clustered’ masses. (E) Quantification showing that single and clustered DNA masses found in *fip-* and Feo co-depleted animals are dramatically larger than those found when *fip-* or Feo are depleted independently. Scale bar: (A) = 10um, (B) = 100um, (C and D) = 50um

## Discussion

FIP was originally named Sip2 for a reported Y2H interaction with Septin 1 and Septin 2 (Shih, Peifer 2002); however, no further studies have reported on Sip2 and our work was not able to confirm this interaction. The highly dynamic, cell cycle-dependent localization of FIP suggested it plays several roles, and our FIP loss-of-function (RNAi in S2 cells and mutant animals) analysis showed slight defects in MT +end growth and an increase in the duration of mitosis. These defects are likely attributed to the localization of FIP to MT +ends in interphase and the perichromosomal region prior to anaphase onset, respectively (Figure 10). In addition, we uncovered a role of FIP in late stages of mitosis where we uncovered phenotypes consistent with cytokinesis defects in *fip-* mutant animals, both in epithelial cells of imaginal wing discs and in the developing larval central nervous system (Figure 10). These data, in combination with FIP::GFP localization at the central spindle following anaphase onset, suggested that FIP does play a role during cytokinesis, but likely through a septin-independent mechanism.

**Figure 10.**
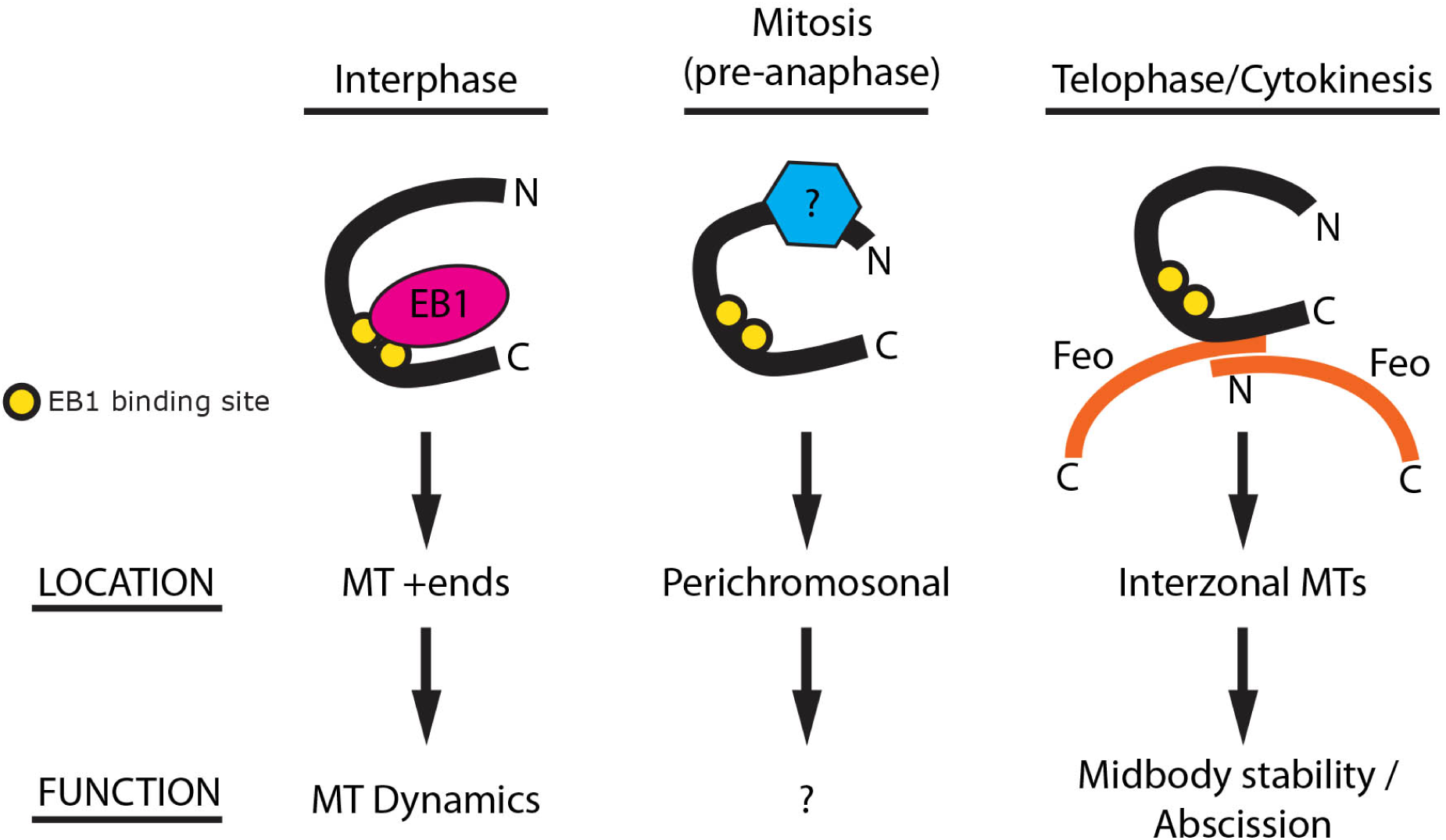
Model of FIP function. FIP is a multifunctional protein with highly variable localizations. In interphase, FIP localizes to MT +ends through EB1 and appears to slightly regulate MT growth. In metaphase, the localization of FIP to the perichomosomal region, the extended mitotic duration, and the higher mitotic index all support an unknown role in mitosis prior to anaphase onset. In later stages of mitosis, the C-terminus of FIP bind the N-terminal region of Feo to regulate interzonal MT stability, ensuring fully accurate cytokinesis and abscission.

Analysis of FIP::GFP shows that it localizes to MTs, and loss of *fip* resulted in the dissipation of interzonal MTs at the end of mitosis three times faster than in controls. This suggests that the stability of the antiparallel interzonal MTs is reduced, and thus, we focused our attention on understanding how FIP regulates MTs in anaphase and telophase. Given that overexpression of FIP did not force its localization onto the MT lattice in interphase or on astral MTs in mitosis, we determined that FIP was likely not a MAP and that FIP regulates interzonal MT stability through additional proteins. Based on our data and published literature, we hypothesized that FIP functions through one of these candidate proteins - EB1, Klp61F (Kinesin-5, Eg5 in mammals), Abnormal Spindle (Asp), and Feo. Using mutant analysis (*fip*^*Δ1-2*^ eliminated EB1 as a candidate), protein localization of FIP::GFP (does not localized to the spindle in metaphase eliminated Klp61F as a candidate), and direct protein-protein interaction via Y2H analysis (an interaction with Feo, but not Asp), we determined that FIP binds MTs via the crosslinking protein Feo.

The next step was to determine if FIP and Feo functioned in the same pathway to regulate late stages of mitosis. Our live dual color imaging revealed an identical localization for FIP and Feo, supporting the Y2H and co-IP data that they function in the same molecular complex. To investigate the phenotype of *feo-* cells, we were unsuccessful in using the null mutant *feo*^*EA86*^ (Perrimon et al., 1989; Verni et al., 2004) as these animals died in early larval stages,highlighting the critical nature of Feo. We therefore utilized RNAi knockdown of Feo, which allowed for the analysis of late third instar larval brains, the exact developmental time where we identified large DNA aggregates (polypoid cells) in *fip-* animals. Analysis of *feo*^*RNAi*^brains also showed large DNA aggregates, very similar to *fip-* mutants. The *fip-* and *feo*^*RNAi*^ double loss of function showed a 4-fold increase in the number of polyploid cells, in addition to the formation of much larger DNA masses and aggregates, nearly 100x the size of normal NB nucleus. Importantly, overexpression of Feo::GFP nearly completely suppressed *fip-* polyploidy, suggesting that FIP falls upstream of Feo in a functional pathway.

The phenotypes we have observed, particularly the rapid dissipation of MTs in *fip-* cells, in combination with our finding that FIP directly binds the N-terminus of Feo, allows us to propose several models. For example, FIP could influence the dimerization state of Feo, which is required for crosslinking MTs (Figure 10). Loss of FIP in this model would lead to a shift toward a monomeric state of Feo, leading to reduced MT crosslinking and effectively reduced interzonal MTs. A second hypothesis is based on previous work showing that Kif4 binds to the N-terminal dimerization and rod domain of PRC1 to recruit MT catastrophe factors and facilitate antiparallel MT ‘pruning’ (Bieling et al., 2010; Nguyen et al.), which maintains proper MT overlap from each spindle pole. Loss of Kif4 leads to increased MT overlap, elongated spindles, and broadening of PRC1 localization in the central spindle (Nguyen et al.; Shrestha et al., 2012; Zhu and Jiang). Loss of FIP gives the opposite result where interzonal MTs rapidly dissipate, possibly due to an increase in MT catastrophe. Thus, one hypothesis is that FIP normally works to maintain the correct amount of MT overlap by limiting the amount of Klp3A (*Drosophila* Kif4) bound to Feo. In this scenario loss of FIP would result in increased Klp3A levels, increased recruitment of catastrophe factors, and elevated interzonal MT pruning. Other hypotheses would include mechanism by which FIP modulates Feo function via competitive or cooperative binding with the many other Feo binding partners. These proposed models represent exciting future directions and will benefit greatly from *in vitro* biochemical studies using purified components. Such experiments will help identify a possible mammalian ortholog of FIP as, while there is no direct sequence ortholog, there is likely a functional ortholog that would regulate PRC1.

## Methods

### Cell culture, transfection, vectors, and dsRNA knockdown

*Drosophila* S2 cells (Invitrogen) were maintained in SF900 media (Gibco) supplemented with Antibiotic-Antimycotic (Gibco) at 25°C. FIP (isoform B) and Feo were PCR amplified and cloned into Entry vectors for the Gateway system (Invitrogen), using the following primers: FIP Forward 5’-CACCATGTCTGGCCTCAAGAAATTCC-3’, FIP Reverse 5’-AAACTTTTGGGCATCTCTTATTGT-3’, Feo Forward 5’-CACCATGAACTCGCCGAGCGCCATTG-3’ and Feo Reverse 5’-GAACTGTCTGCGCGGCTGCACG-3’. We used the manufacturer’s protocol in combination with Gateway destination vectors from the *Drosophila* Gateway Vector Collection, and our personal collection, to generate fusion constructs tagged with GFP, Flag, mNeonGreen, TagRFP, or Halo under the control of the Actin5c promoter. The following constructs were generated: FIP::GFP, GFP::FIP, TagRFP::FIP, mNeonGreen::Feo, Flag::FIP, GFP::Feo, and GFP::EB1. Transfection was performed as described previously (Schoborg et al., 2015) with the following differences: 1μg of vector was used for 1-5 million cells. For dsRNA treatment, ∼5 million cells were collected two to three days after passaging and resuspended with 2ml of fresh SF900 media with 20µg dsRNA. Every two days, media was replaced with fresh media and dsRNA. Cells were either fixed and stained on day 6 or transfected on day 4 and allowed to grow for two additional days before imaging. An identically treated well lacking dsRNA was included to control for growth conditions. The following primers sequences were obtained from the Harvard *Drosophila* RNAi Screening Center and used to generate templates for T7 RNA synthesis reactions (Promega) from FIP cDNA: 5’-ATGGCTGCAAAAAGGCTAC-3’, 5’-CTAAACGTCGACGAATTGTT-3’.

### Staining and immunofluorescence

S2 cells were allowed to adhere to Con-A coated coverslips for 20-30 minutes, briefly washed with PBS, and then fixed with either pre-chilled anhydrous methanol for 20 minutes at - 20°C or 4% paraformaldehyde (PFA, Electron Microscopy Sciences) in PBS for 20 minutes at room temperature. If necessary, cells were counterstained for one hour with DAPI and Alexa 568 conjugated phalloidin (Invitrogen), rinsed 3x and then mounted in Vectashield (Vector laboratories). Phalloidin was desiccated in a glass well and resuspended in PBS before using to remove methanol. Larval brains and wing discs were obtained from wandering 3rd instar larvae and fixed as inverted carcasses in 4% PFA diluted in PBS with 0.05% Triton X-100 for 30 minutes at room temperature while rotating. The only exception to this protocol was used to prepare the wing discs in Figure 7 which were fixed in cold PBST with 4% PFA. One explanation might be that the chilled wing discs might have better exposed the Pnut antigen as the anti-Pnut antibody did not work otherwise. All tissues were dissected in Schneider’s *Drosophila* Medium (Gibco) supplemented with Antibiotic-Antimycotic (Gibco). Tissues were blocked with 5% normal goat serum (NGS) in PBSTx (PBS with 0.05% or 0.3% Triton-X 100) for one hour at room temperature and incubated with primary antibodies diluted in PBSTx + 5% NGS overnight at 4°C. Samples were washed 3X in PBSTx for 20-30 minutes to an hour at room temperature then incubated with secondary antibodies diluted in PBSTx + 5% NGS with DAPI and phalloidin for 2-24 hours at room temperature or 4°C. Tissues were washed 3X in PBSTx then mounted in Vectashield. The following primary antibodies were used: mouse anti-Pnut 1:200 (4C9H4 DSHB), mouse anti-lamin 1:100 (ADL84.12 DSHB), mouse anti-GFP (JL-8) 1:5000 (Clonetech), mouse anti-flag 1:10,000 (M2 Sigma-Aldrich), rabbit anti-GFP 1:1000 (ab 290 AbCam), rabbit anti-phosphorylated histone H3 1:1000 (Millipore). Goat secondary antibodies raised against the appropriate species were Alexa Fluor 488, 568, or 647 conjugated 1:500 (Invitrogen). For live imaging of Halo-tagging proteins, cells were stained with the JF549 ligand for >1hr prior to live imaging. A 1mM stock solution of JF549 in DMSO was added to culture media at 1:200 for a final working concentration 5nM of ligand. For fixed imaging of Halo-tagged proteins, S2 cells expressing GFP-Tub and Ht-Sip2 were incubated for 1hr with 100nM TMR-HL, washed several times with PBS, fixed for 10 minutes in cold MeOH, rehydrated in PBSTween, and stained an hour with DAPI prior to mounting.

### Immunoprecipitation and Immunoblotting

For IPs, 4 ml of transfected *Drosophila* S2 cells (either treated with 30uM colchicine for 16 hours to increase the mitotic index to ∼20% or with ethanol as a control), were spun down, had the supernatant removed, and then resuspended in 1mL of RIPA Buffer (10 mM Tris-HCl, pH 7.5, 140 mM NaCl, 1 mM EDTA, 0.5 mM EGTA, 1% Triton X-100, 0.1% sodium deoxycholate, 0.1% sodium dodecyl sulfate, and 0.25μM protease inhibitors (Pierce Protease inhibitor mini EDTA Free (Thermo Scientific)). 50μL of the lysate was removed and saved as the “input” sample while the rest of the lysate was combined with Protein-A-conjugated dynabeads (Life Technologies) prebound to antibody (see below) and incubated for 2 hours at 4°C, rotating. The supernatant of the lysate and bead mixture was removed and the beads were washed 3 times in lysis buffer. The beads were moved to a fresh tube during the final wash, eluted with 30μL of Laemmli buffer, and boiled for 5 minutes. For antibody binding, 50μL of dynabeads and 1μL of rabbit anti-gfp antibody (ab 290 AbCam) were added to 1mL of PBST (PBS and 0.01% Tween-20). The bead mixture was incubated at 4°C, rotating, for 1-2 hours. The elute was removed and resolved on SDS-PAGE along with the input samples and transferred onto nitrocellulose membrane. Western blots were stained with primary antibodies ((mouse anti-GFP (JL-8) 1:5000 (Clontech) or mouse anti-flag 1:10,000 (M2 Sigma-Aldrich)) and anti-mouse peroxidase conjugated secondary antibodies 1:5000 (Thermo Scientific). Western blots were visualized using SuperSignal West Dura Extended Substrate (Life Technologies) and imaged using a ChemiDoc MP Imaging System (BioRad).

### Fly stocks and genetics

The following fly strains were used: Cyo,*P{GFPnls},* Cyo,*P{2xTb*^*1*^*-RFP},* TM6B,*Tb*^*1*^, *Antp*^*Hu*^andTM6C,*Sb*^*1*^,*Tb*^*1*^balancerchromosomes, *Tub-GAL-4* (Bloomington stock (BS) #5138), *sqh-GAL4* and *ubi-Moe::GFP/CyO; TM3/TM6* (gift from Dan Kiehart), *Dpn-GAL4* (gift from Makato Sato), *UAS-EB1-GFP* (gift from S. Rogers and B. Eaton), *G147* (Morin et al.), *H2AV-mRFP* (BS #23650), *Df(2L)Exel7029* (BS #7802, Sep2::GFP (BS #26257), *Ubi-p63E-feo::GFP* (BS #59274), *Ubi-p63E-feo::mCherry* (BS #59278), *UAS-Feo*^*RNAi (HM)*^ (BS #28926), *UAS-Feo*^*RNAi (GL)*^ (BS #35467). All fly stocks were maintained on standard cornmeal-agar media at room temperature (19-23°C), experimental crosses were kept at 25°C until fixation or live imaging. Figures 8D and E were obtained from homozygous *fip*^*7*^ larvae, in all other cases the genotype *fip-* refers to *fip*^*7*^/*Df(2L)Exel7029*.

### CRISPR and transgenic animals

Two CRISPR guide RNA sites were determined using http://flycrispr.molbio.wisc.edu/ 5’-GAATACTATTGCCAGAAGGT-3’ and 5’-GCGACGCTGAGGAATACCAG-3’. Each guide was cloned into a separate pU6-Bbsl plasmid and equimolar amounts were injected into Cas9 embryos by BestGene Inc Chino Hills, CA). Injected males were mated to *yw* females and progeny were screened for non-homologous end joining (NHEJ), thus excising ∼80% of the FIP locus, by single wing PCR using the primers 5’-GCAAAGGCGCGTCGATCGTTGGC-3’ and 5’-TAGCGGAGCAGTACCAGACTTCTGGG-3’. We isolated 7 clones and sequenced the FIP locus of each, revealing 5 different alleles, each of which differed slightly in the exact NHEJ product. We found no significant difference in the penetrance/severity of our primary phenotype (polyploid neuroblasts in the larval brain). Therefore, we randomly selected FIP^7^ for use in this study.

### Microscopy and live cell imaging

Live and fixed cell imaging was performed on a Nikon Eclipse Ti inverted microscope equipped with a CSU22 spinning disk unit (Yokogawa) and an ORCA-Flash4.0 CMOS camera (Hamamatsu) with either a 40x/1.30 NA plan Fluor objective, sometimes in conjunction with a 1.5X tube lens, or 100x/1.49 NA TIRF objective. Solid state laser lines at 405, 491, 561, and 642 nm (VisiTech International). Microscope control and image acquisition were performed using Metamorph (version 7.8.13.0).

Live cell imaging in Figures S1A, B, S2A, and fixed imaging in Figure S3C were performed in super-resolution mode on a Zeiss LSM 880 equipped with an Airyscan module using a 63x/1.4 NA objective. Figures 4A-C, S3B, and S4A were collected using Airyscan and a Fast module set for 0.5x Nyquist sampling on a 40x/1.4NA Plan-Apochromat objective. All image acquisition was performed using Zen Black software (version 2.3), where Airyscan technology was utilized, raw data were processed and deconvolved within Zen, processing strength was automatically computed within Zen.

For live imaging of S2 cells, 200µl of resuspended cells were plated onto a glass-bottom 35mm dish (MatTek) coated with 15µg of Concanavalin A (Con A) and allowed to settle for 20-30 minutes before imaging. For extended time-lapse imaging, 1.5ml of conditioned SF900 media was added to the dish, instead of fresh Schneider’s medium, after cells had adequately adhered. When imaging cytokinesis in S2 cells, uncoated dishes were used, as Con A inhibits cytokinesis, and cells were imaged immediately. In Figure 2B, Hoechst 33342 and SiR-Tubulin (Cytoskeleton) were used at 1.5µM and 0.1µM respectively.

Larval brains and wing discs were dissected from wandering 3rd instar larvae in Schneider’s *Drosophila* Medium (Gibco) supplemented with Antibiotic-Antimycotic (Gibco). Brains and wing discs were briefly washed into PBS and then gently adhered to a poly-L Lysine coated glass-bottom 35mm dish in PBS before replacing PBS with 1.5ml Schneider’s medium. Wing discs were mounted with the peripodial epithelium facing the coverslip, while brains were mounted with the ventral surface facing the coverslip. In all cases, we used #1.5 coverslips.

### Image processing and analysis

In all cases where measurements were inherently subjective, images and image series were randomly blinded before analysis to avoid cognitive bias. In all cases where live measurements were compared between multiple conditions, all imaging was performed on the same day to avoid changes in laser level or ambient temperature which could influence fluorescence or cell physiology.

All image processing, quantification, and false coloring was performed using FIJI (Image J, NIH, Bethesda, MD). Image quantification was performed exclusively on raw data. If rotation was necessary (e.g. to generate kymographs or to reorient tissues), images were rotated only once and bicubic interpolation was used. Unless disclosed here, image manipulation was limited to linear contrast stretching to best represent the main message of each image. Figure 1B and S4E were filtered of outlying (hot) pixels using the “remove outliers” command, with a threshold of 75 gray values (∼50% of the true dynamic range). Images in Figure S1B were rotated so that each tip is oriented with the + end facing right.

All image series used for temporal-color coding were first corrected for bleaching using histogram matching so that truly static pixels present as grey in the resulting color-coded image and not a reddish yellow. Color coded projections were generated using the Temporal-Color Code feature in FIJI. Images in Figure 5 were filtered with a gamma correction (gamma = 0.65 for phalloidin and 0.75 for DNA) to better represent a mix of bright and dim signals, and rotated so that the anterior of the brain faces up.

Kymographs were generated by rotating an image series so that movements of interest occurred predominantly along the x-axis. A rectangular region encompassing the region of interest was resliced without interpolation and maximum intensity projected so that each frame of the original image series represents a single line of pixels on the resulting kymograph.

Enrichment measurements of FIP at MT tips and spindle midzones, and Moesin in the nascent cytokinetic furrow, were calculated by manually defining a region of interest (microtubule tip, nascent furrow, or spindle midzone), and a control region adjacent to the region of interest. Enrichment was calculated using the following equation:

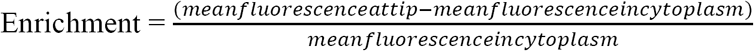

Microtubule growth rates were calculated using a manual tracking plugin for Fiji, each data point represents the average speed of a single MT. Multinucleate, metaphase, and anaphase frequency in FIP dsRNA treated S2 cells were calculated by dividing the number of multinucleate, metaphase, and anaphase cells by the area of a single field of view (∼37,000 µm^2^). Micronuclei and binucleate frequency were calculated by dividing the number of aberrant cells in a single wing disc epithelium by the total number of scoreable cells (i.e. those easily represented by one or two focal planes) in the region.

Nuclear envelope breakdown (NEB) to anaphase timing was determined based on a sudden change in the nuclear outline and the first detectable separation of sister chromatids respectively. Cytokinesis timing was calculated by counting the time between the formation of the nascent furrow and the last frame where a clear separation between furrows could be resolved.

Wing disc size and PH3+ cells were measured from third instar wandering larvae, 96 hours after egg laying. Wing discs were dissected, fixed, and stained with DAPI. The outline tool in Fiji/ImageJ was used to manually encircle the wing disc using the DAPI channel as a reference. The number of PH3 positive cells per wing disc was determined using thresholding in Fiji. Proper thresholding was determined by comparing PH3 counts from Fiji to those initially counted manually to ensure that the automated thresholding values were correct.

Maximum sister chromosome separation was determined by measuring the space between H2AV masses at their most distant point (153 ± 33 seconds after anaphase in mutants, 195 ± 17 seconds in controls). Midbody duration was scored by calculating the time from anaphase onset until the GFP::Jupiter, GFP::Sep2, or GFP::Moesin fluorescence at the midbody was no longer detectable above the background cytoplasmic fluorescence. Only midbodies that persisted within the central slices of the Z stack were used for analysis to avoid midbodies that simply drifted out of the focal plane.

Polyploid cells in the larval CNS were scored based on intense DAPI staining, a larger than average nucleus, abnormal DNA architecture, and multiple nuclei (determined by lamin staining) contained within the same cell. For quantifications, polyploid cells were counted as “one” even if they contained multiple fragmented nuclei or multiple intact nuclei within the same cell as determined by actin staining. For the large DNA aggregates reported for the *fip-* and Feo co-depleted animals, those polyploid cells were also quantified as one polyploid cell although it is hypothesized that multiple polyploid neuroblasts may have fused together to form the large DNA aggregates based on the dramatically larger cell size and DNA content within those neuroblasts. Polyploid nuclei of the double mutants were considered large single or large clustered masses if the cross-sectional area of the DNA exceeded 1000 um^2^.

### Yeast-Two-Hybrid analysis

FIP, Feo, Asp, Sep1 and Sep2 full-length and FIP and Asp pieces were amplified from cDNA clones by PCR using Phusion (Thermo Fisher Scientific, Waltham, MA) and with the primers shown in Table 1. PCR products were then introduced into Gateway Entry vectors using the pENTr/D-TOPO Kit (Thermo Fisher Scientific). All Y2H experiments were then conducted as described previously in (Galletta and Rusan, 2015). In brief, FIP, Feo, Asp, Sep1, and Sep2 full-length and FIP, Feo, and Asp pieces were introduced into pDEST-pGADT7 and pDEST-pGBKT7 using Gateway technology (Thermo Fisher Scientific), transformed into Y187 or Y2HGold yeast strains (Takara Bio USA, Mountain View, CA), and grown in −leu or −trp media. After mating bait and prey strains, diplods containing both were selected on −leu, −trp(DDO) plates, then replica plated onto plates of increasing stringency: DDO; −ade, −leu, −trp,−ura (QDO); −leu, −trp plates supplemented with Aureobasidin A (Takara Bio USA) and X-α-Gal (Gold Biotechnology, St. Louis, MO) (DDOXA); and −ade, −leu, −trp, −ura plates supplemented with Aureobasidin A and X-α-Gal (QDOXA). Interactions were scored based on growth and the development of blue color as appropriate. All plasmids were tested for the ability to drive reporter activity in the presence of an empty vector (autoactivation). Plasmids that conferred autoactivity were omitted from further analysis.

## Acknowledgements

We thank Brian Galletta and Matthew Hannaford for critical reading and discussion of the manuscript, and Ryan O’Neill for significant help with fly genetics. We thank Xufeng Wu and Chris Combs at the NHLBI Light Microscopy core facility for training, guidance, and discussion. This work is supported by the Division of Intramural Research at the National Institutes of Health/National Heart, Lung, and Blood Institute (1ZIAHL006126 to NMR).

## Supplemental Figure legends

**Figure S1.**
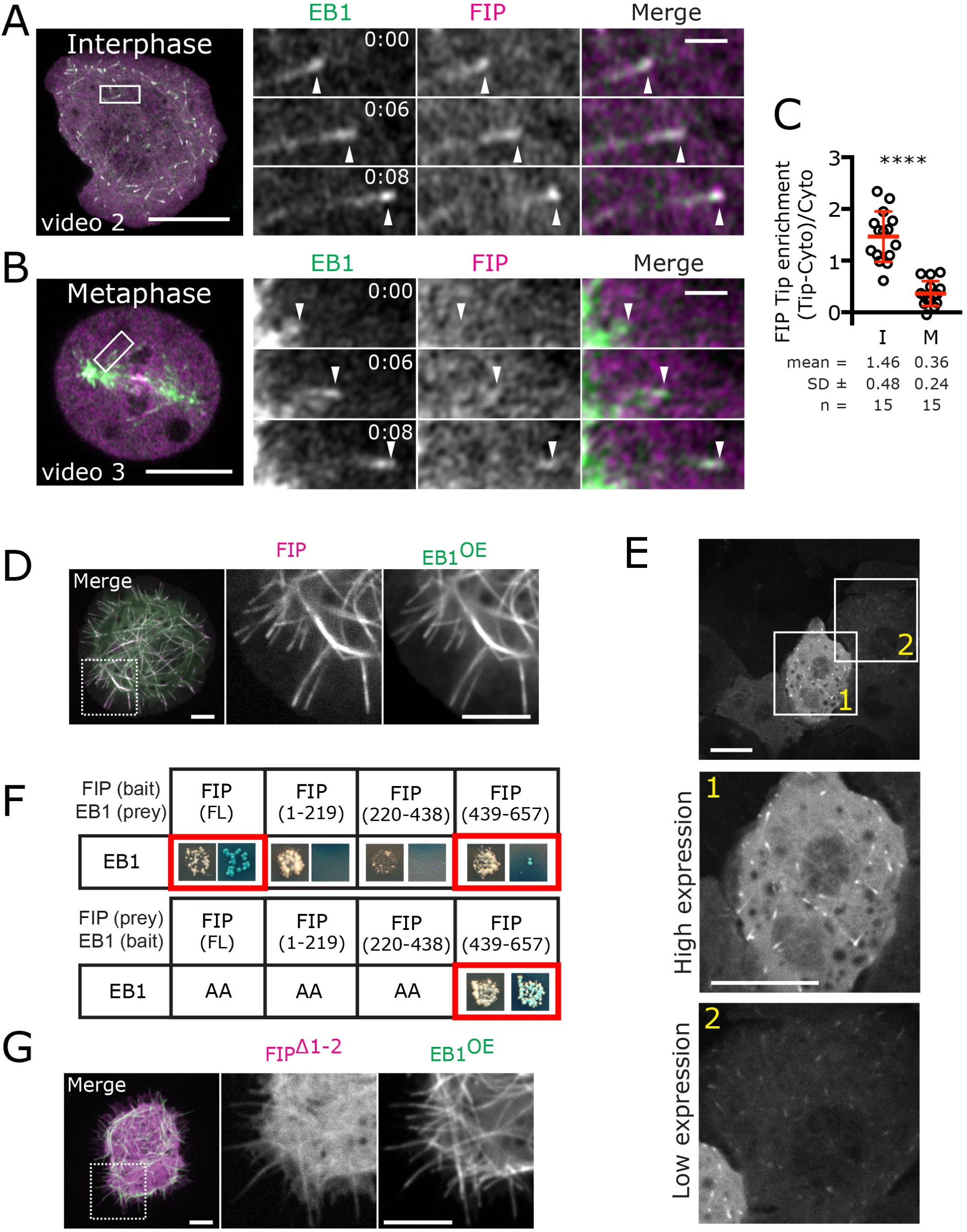
FIP directly binds EB1 and tracks MT +ends during interphase. (A, video 2) interphase and (B, video 3) metaphase S2 cells coexpressing Ht::FIP (JF549 ligand, magenta) and EB1::mNG (green). Boxed regions are enlarged to the right to indicate colocalization of EB1 comets with FIP at a MT +end. (C) Quantification of FIP enrichment at MT +ends during interphase (I, 3 cells) and metaphase (M, 3 cells); each point represents a single MT +end measurement (**** = P < 0.0001). (D) S2 cells grossly overexpressing EB1 (EB1^OE^, green) decorate the entire MT lattice and recruit FIP (magenta). (E) Single image from a time-series of S2 cells transfected with GFP::FIP showing a high-(#1) and low-expressing (#2) cell illustrating that, regardless of expression level, FIP is not forced onto the MT lattice. (F) Y2H analysis showing that only FIP^FL^ and FIP^439-657^ directly bind EB1 (Blue colonies). AA = auto activator for the Y2H system, thus no interaction information is gained. (G) EB1^OE^ (green) is not able to recruit FIPΔ^1-2^ (magenta) to the MT lattice. Scale bars: (A, B, D, G) = 5µm, (A, B enlarged images) = 1µm, (E) = 10 µm.

**Figure S2.**
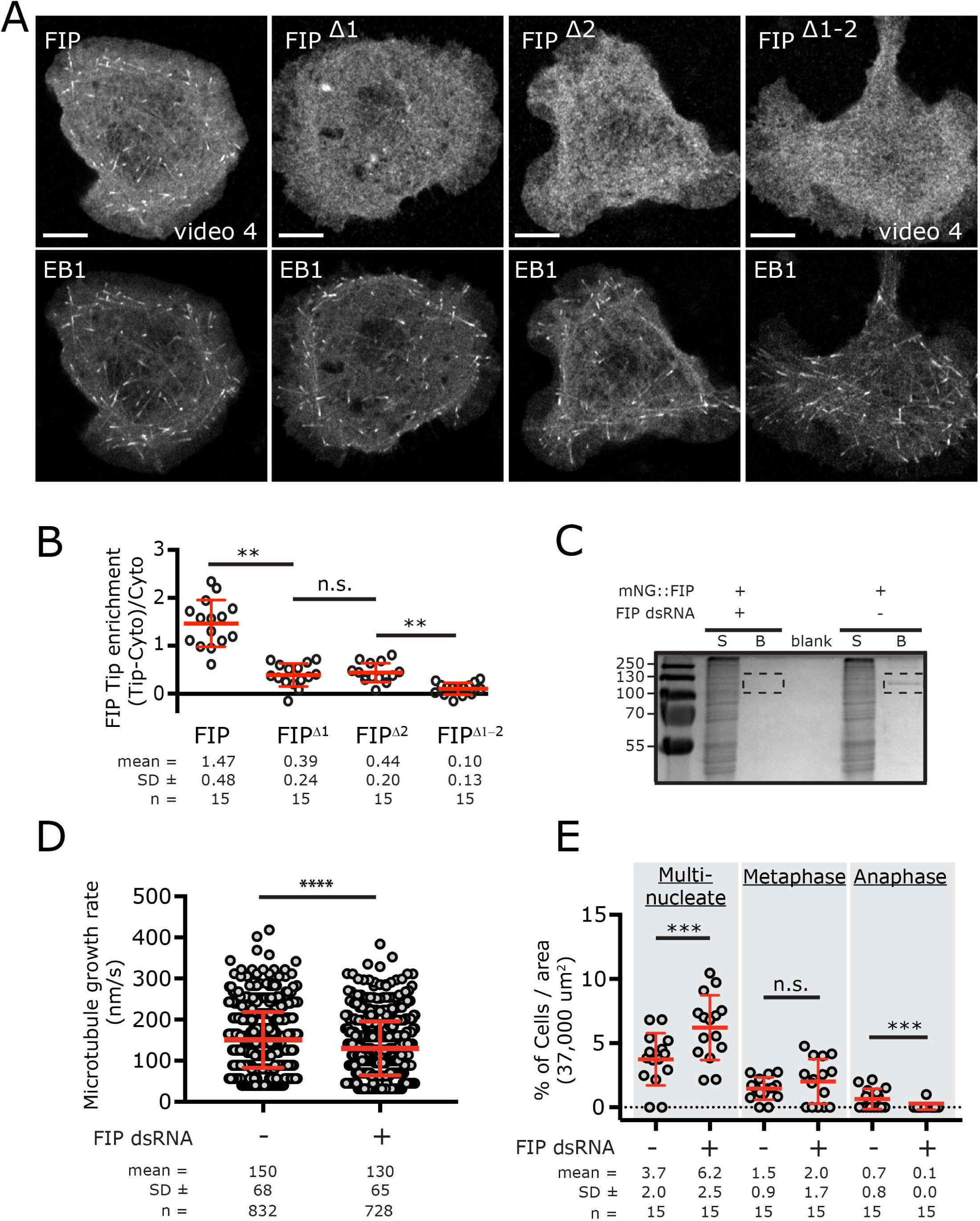
FIP is required for efficient MT growth in interphase and efficient cell division. (A) Wild-type FIP, and SxIP motifs mutants FIP^Δ1^, FIP^Δ2^ and FIP^Δ1-2^were HaloTagged (JF549 ligand) and coexpressed with low levels of EB1::mNG (video 4). (B) Quantification of fluorescence enrichment of FIP variants at the MT +end over cytoplasmic levels; each point represents a single MT +end measurement (15 measurements from 3 cells; ** =P < 0.01 (C) Polyacrylamide gel stained with coomassie brilliant blue showing a failure to immuno-precipitate mNG::FIP from cells treated with dsRNA against FIP, indicating successful dsRNA knockdown. Supernatant from each condition are run in the “S” lane, protein eluted from anti-mNG nanobody beads are run in the “B” lane. (D) Quantification of MT growth rates showing a slight but significant decrease in growth velocity in FIP depleted S2 cells; each point represents the average velocity of a single MT track (**** = P < 0.0001). (E) Quantification of cell cycle stages and mitotic defects in wild-type and dsRNA knockdown S2 cells (*** = P < 0.001). Scale bars: (A) = 5µm.

**Figure S3.**
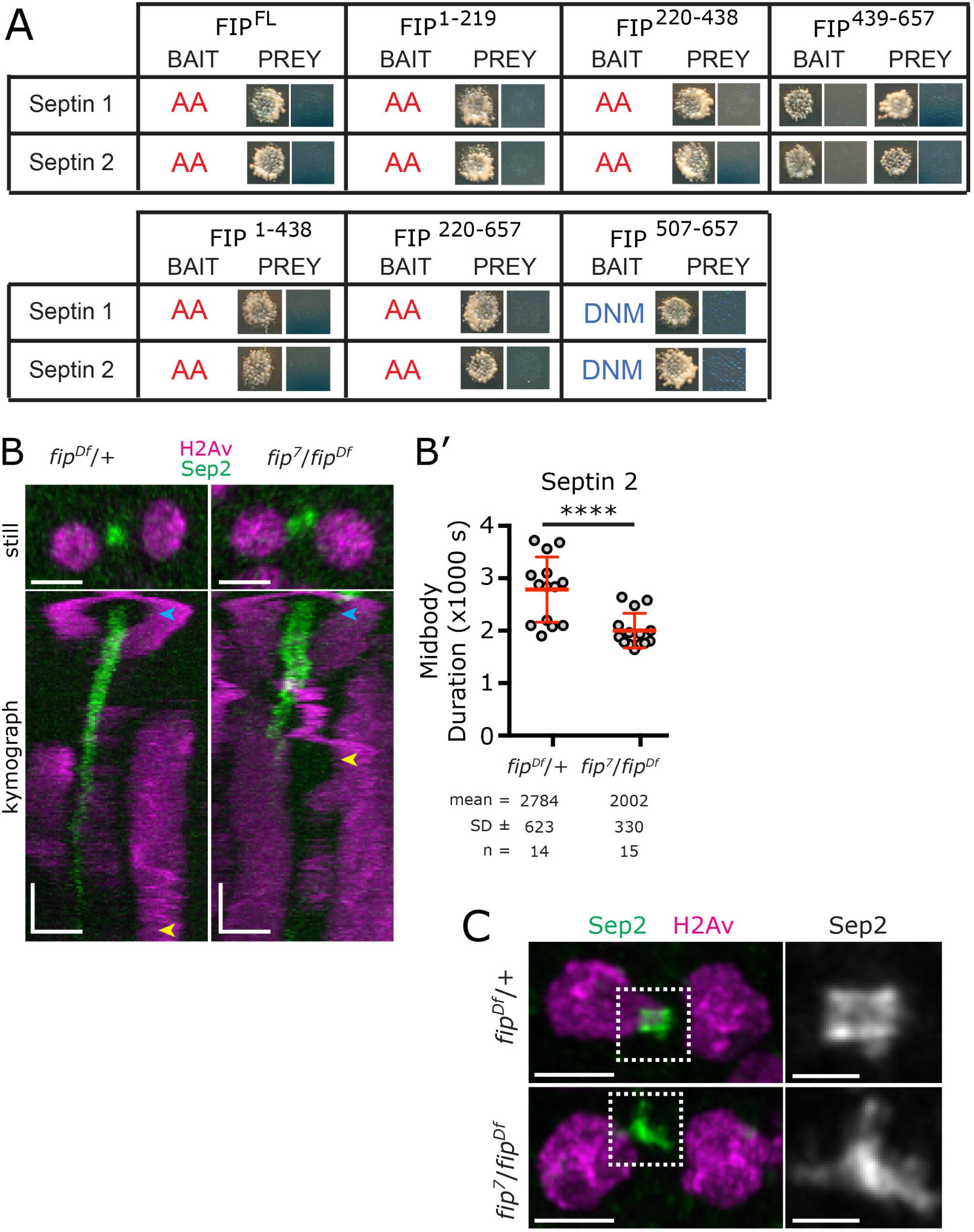
FIP indirectly helps organize the Septin network. (A) Colonies from Y2H analysis between FIP and Septin 1 or Septin 2. No blue colony growth indicated no detectable interaction. AA = auto activator for the Y2H system, thus no interaction can be tested. DNM = did not mate (B) Stills and kymographs from telophase *fip-* and control wing discs expressing Sep2::GFP and H2AV::mRFP (marking chromosomes). Kymographs below each still begins at anaphase (lateral separation of the chromosomes, magenta streaks). Green vertical streak represents midbody formation, note the rapid dissolution of the green signal in the *fip-* kymograph. Blue arrows indicate the frame of the kymograph used to create the still image above, yellow arrows indicate when Sep2::GFP signal fully dissipates. (B’) Quantification of Sep2::GFP signal duration (anaphase to the last frame where Sep2 midbody localization is detectable) showing that *fip-* midbodies lose septins faster than controls; each point represents a single cell. Three wing discs were used for each condition (**** = *P* < 0.0001). (C) High resolution Airyscan images from fixed *fip-* and control wing discs at comparable stages to the blue arrows in (B), Sep2 signal is enlarged in the grayscale images to the right. Horizontal scale bars in (B), = 3µm, vertical scale bars in (B) = 480 seconds. Scale bar in (C) = 3µm, inset = 1µm.

**Figure S4.**
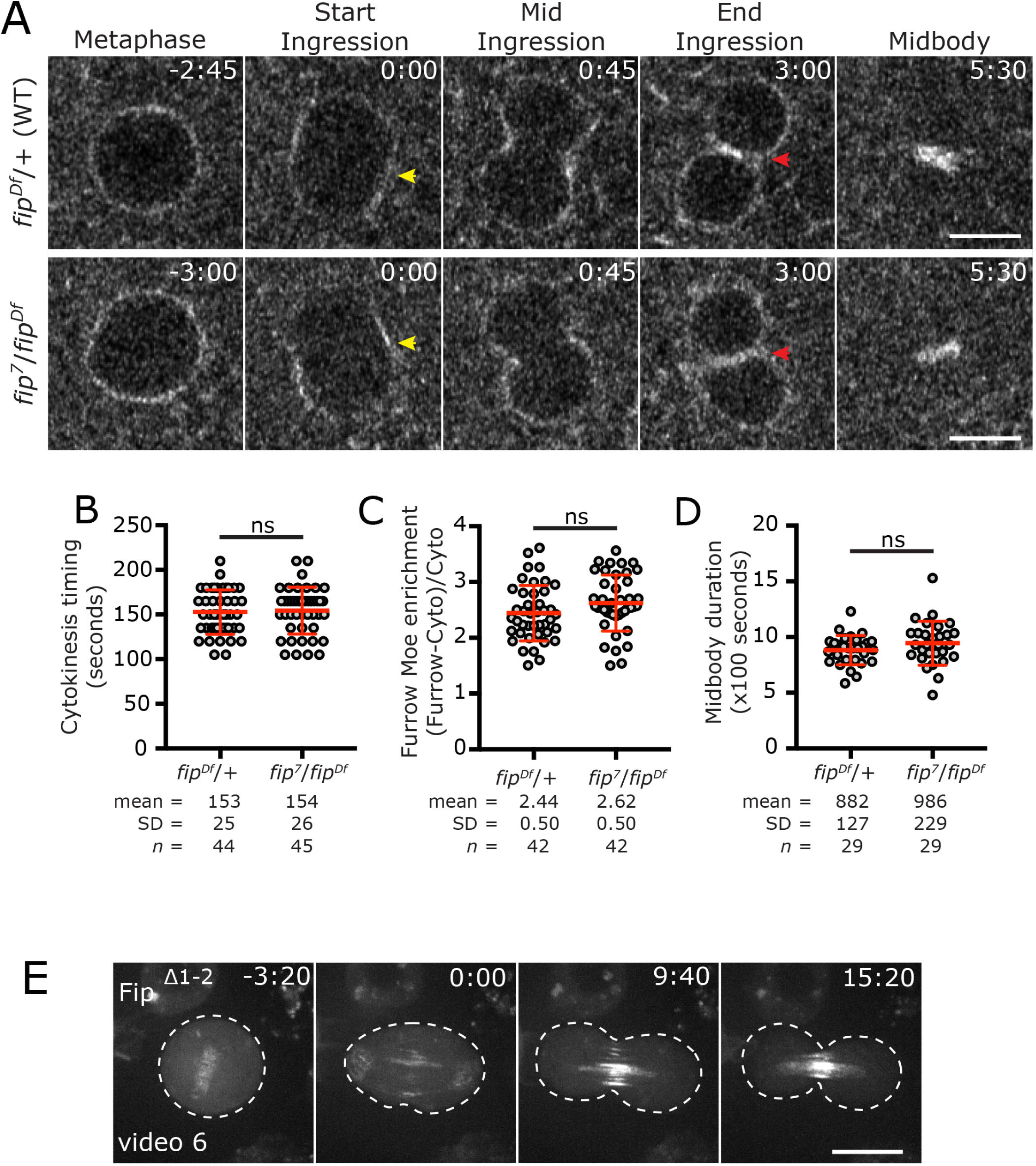
Loss of FIP has no detectable effect on the actin contractile ring. (A) Wing disc cells from animals expressing GFP::Moesin. Dividing *fip-* cells show comparable timing of furrow initiation (yellow arrow) and complete formation of the constricting cytokinetic ring (red arrows). (B, C, D) Quantification of videos as in A for both wild-type and *fip-* cells. In all cases, there is no significant difference. Scale bars in (A) = 5um. Time is in min:sec relative to anaphase onset at 0:00. (E) Representative time series of a S2 cells expressing FIP^Δ1-2^ (video6) from metaphase to late telophase. Time is in min:sec relative to anaphase onset. Scale = 10µm.

**Figure S5.**
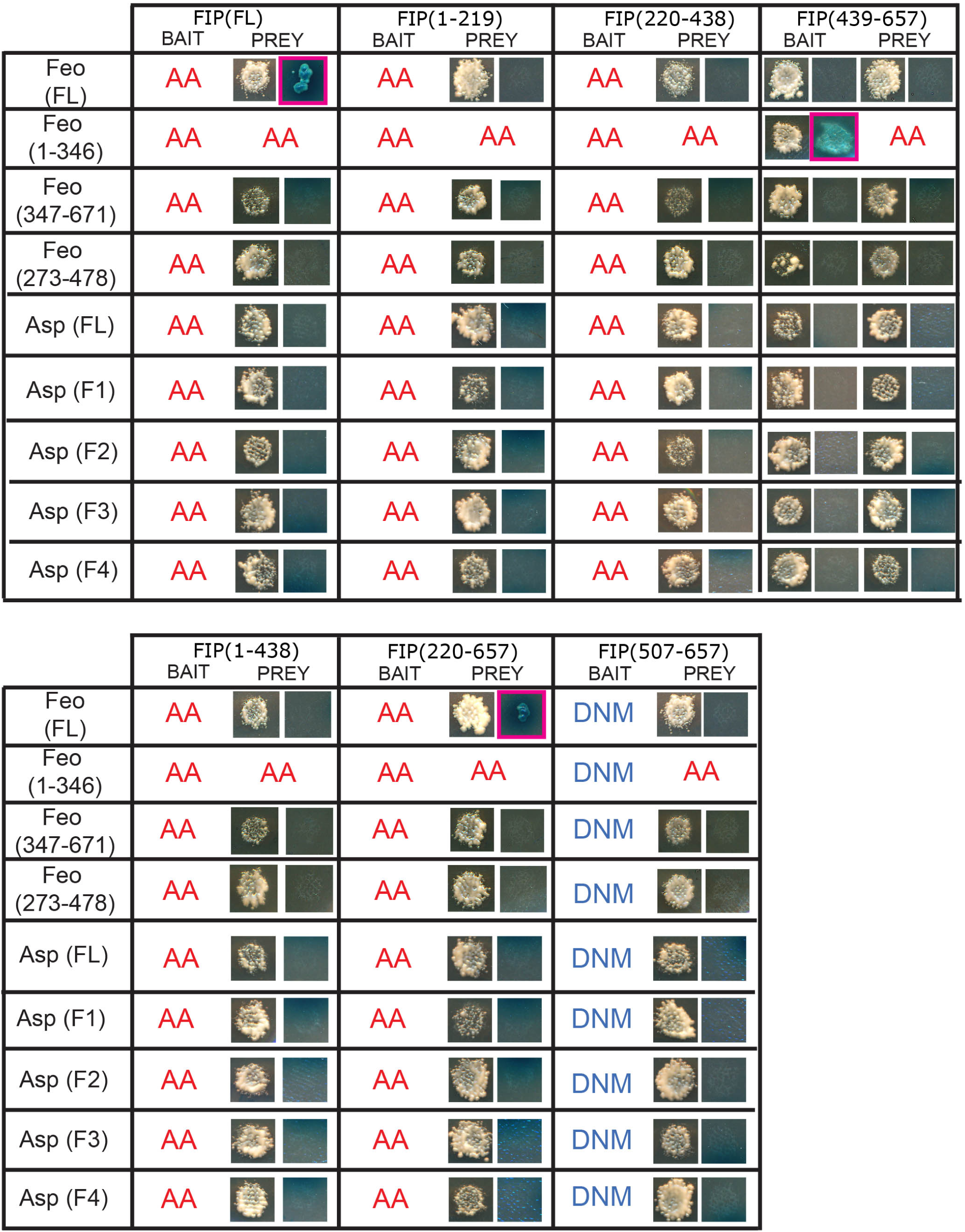
FIP binds Feo, but not Asp. Colonies from Y2H analysis between FIP fragments and each of Asp and Feo. Blue colony growth indicates interaction (pink box) only with Feo. AA = auto activator for the Y2H system, DNM = did not mate, no interaction information can be determined for AA or DNM.

**Figure S6.**
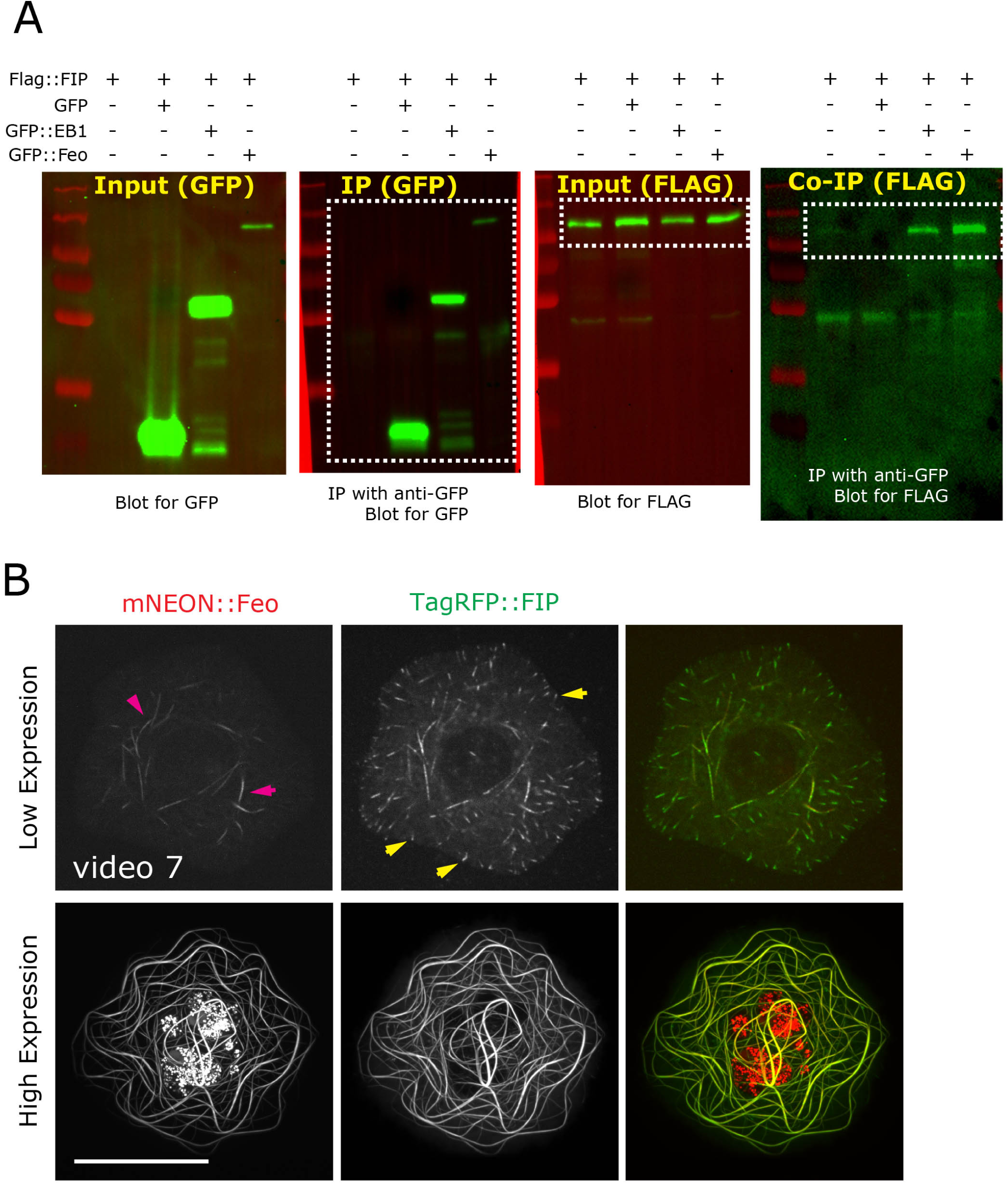
Feo binds and recruits FIP to MTs. (A) Raw data of the blots used to assemble Figure 6B. (B) mNeonGreen::Feo (red) and TagRFP::FIP (green) were co-transfected in S2 cells. Interphase cells expressing low amounts of mNeonGreen::Feo show normal FIP MT +end tracking (yellow arrows) and a small amount of MT bundling (pink arrows). High expressing cells illustrate Feo’s ability to recruit FIP to the MT lattice. Scale bar = 10µm.

**Figure S7.**
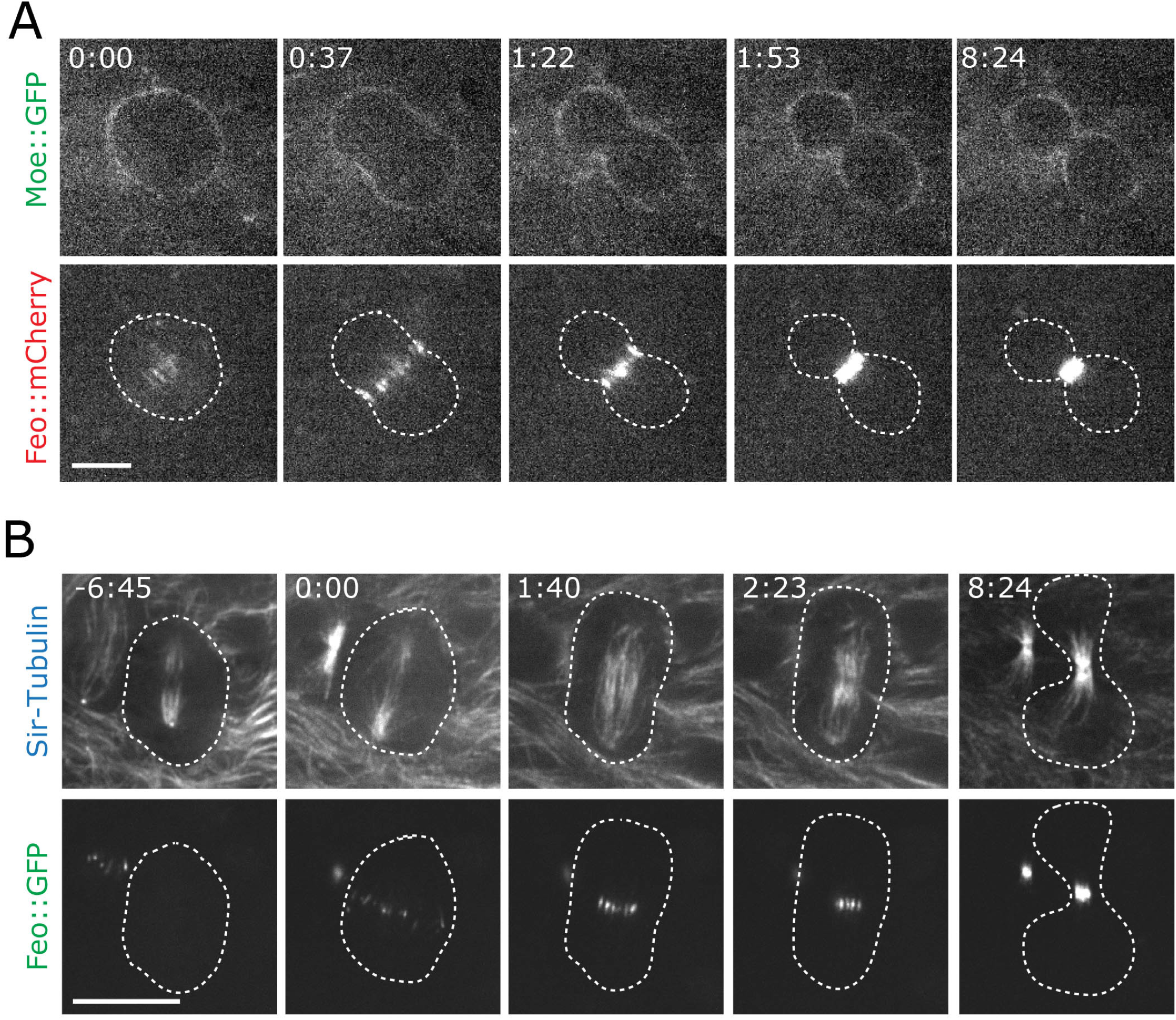
Feo localization in larval wing disc cells. (A) Wing disc cell from animals expressing Feo::mCherry and Moe::GFP. (B) Wing disc cell from animals expressing Feo::GFP and labeled with Sir-Tubulin to image MTs. Both A and B show that Feo localizes to interzonal MTs during cytokinesis as predicted from published data. Scale bars in (A) (B) = 5um. Time is in min:sec relative to anaphase onset at 0:00.

**Figure S8.**
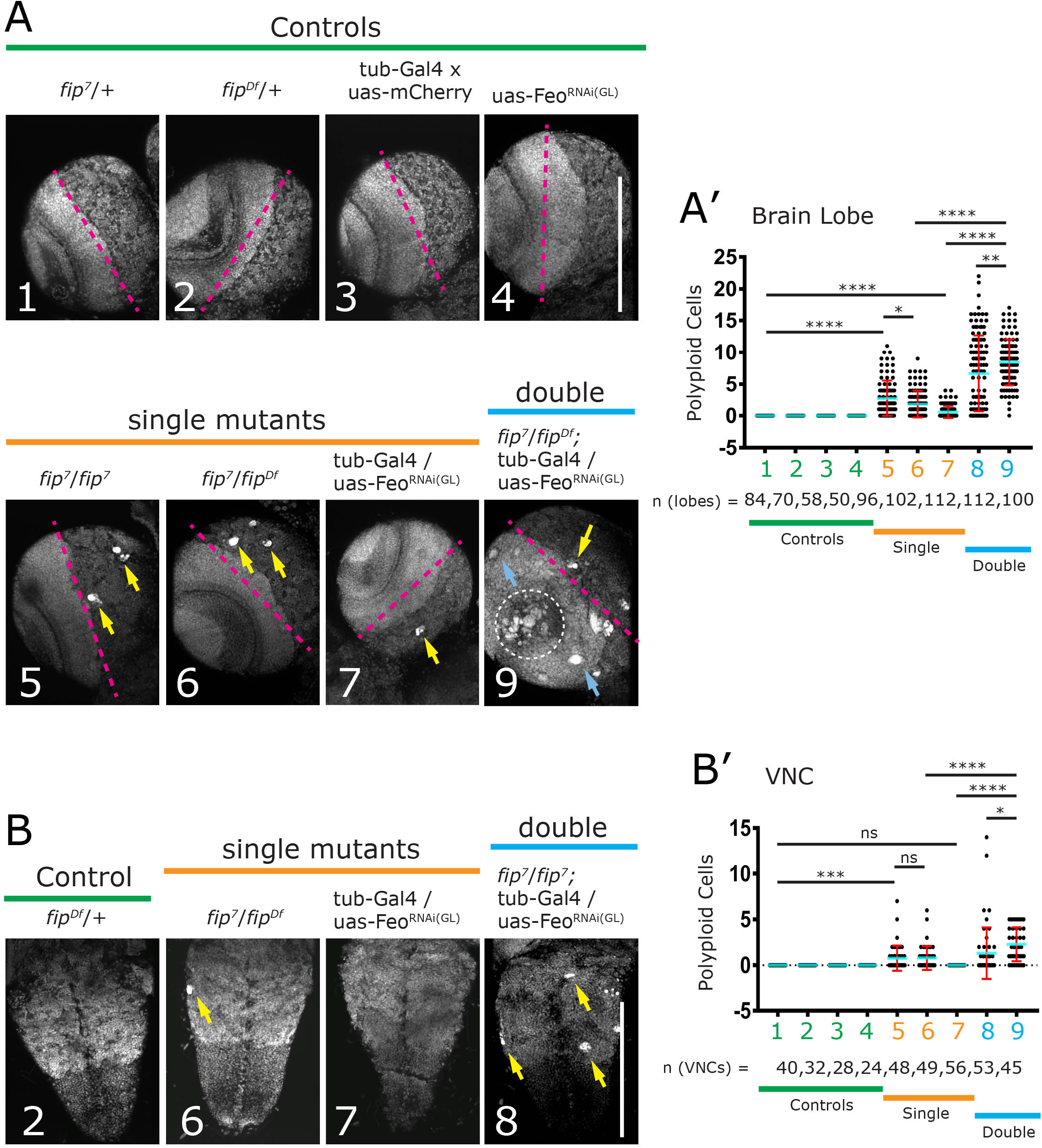
Fip and Feo depletion results in polyploid neuroblasts in the larval brain. Expanded version of Figure 8 with all controls and genotypes. (A) Third instar larval brains stained for DNA (DAPI) in control (green), single mutant (orange), and double mutant (blue) conditions. Pink dotted lines delineate the central brain (right) from the optic lobe (left). Polyploid cells in the central brain (yellow arrows) are present in single mutants while polyploid cells are present in both the central brain (yellow arrows) and optic lobes (blue arrows) in double mutants. (A’) Quantification of polyploid cells per brain lobe. (B) Ventral nerve chord (VNC) stained for DNA (DAPI). Genotypes as indicated above. (B’) Quantification of polyploid cells in the VNC. Scale bar = 150um

## Video List

**Video 1. | FIP is perichromosomal in metaphase and at spindle midzone MTs during anaphase and telophase.** Corresponds to Figure 1B. Cultured S2 cell transiently expressing GFP::FIP. Video begins in metaphase where FIP is concentrated around mitotic chromatin, and extends through late telophase where FIP concentrates around the cell midbody. Image series is maximum intensity projection of six 0.8µm slices/frame, acquired at 20 s/frame and displayed at 10 fps, time is in minutes:seconds relative to NEB, scale bar = 10µm.

**Video 2. | FIP localizes to growing interphase MT +ends.** Corresponds to Figure S1A. Interphase S2 cell coexpressing Ht::FIP (JF549 ligand, magenta) and EB1::mNG (green) illustrating colocalization of EB1 comets with FIP at MT +ends. Boxed frames in upper right show the EB1 channel in greyscale (top), FIP channel in greyscale (middle) and merge (bottom). Image series is a single slice, acquired at 2 s/frame and displayed at 5fps, time is in minutes:seconds, scale bar = 5µm.

**Video 3. | FIP localization to MT +ends approaches background levels in mitosis.** Corresponds to Figure S1B. Metaphase S2 cell coexpressing Ht::FIP (JF549 ligand, magenta) and EB1::mNG (green) illustrating colocalization of EB1 comets with FIP at MT +ends. Boxed frames on right show the FIP channel in greyscale (bottom), EB1 channel in greyscale (middle), and merge (top). Image series is a single slice, acquired at 2 s/frame and displayed at 5fps, time is in minutes:seconds, scale bar = 5µm.

**Video 4. | FIP localization to interphase MT +ends is EB1-dependent.** Corresponds to Figure S2A. Interphase S2 cell coexpressing Ht::FIP (top left), or Ht::FIP^Δ1-2^ (top right), and EB1::mNG (bottom row), illustrating that the FIP^Δ1-2^ is incapable of MT +end tracking, despite clear EB1 MT +end tracking. In both cases, Halo-tagged FIP constructs are covalently linked to a JF549 Halo Tag ligand. Image series is a single slice, acquired at 2 s/frame and displayed at 5fps, time is in minutes:seconds, scale bar = 5µm.

**Video 5. | Loss of FIP leads to failed cytokinesis and increased ploidy.** Corresponds to Figure 5E. Polyploid neuroblast from a *fip-* 3^rd^ instar larval brain expressing GFP::Moesin (magenta) and H2AV::mRFP (green). Neuroblast enters anaphase and attempts to complete cytokinesis, but fails. The contractile ring regresses and the DNA from the would-be daughter cells collapses into a single DNA mass. Frames were captured at different rates, but displayed at 21 fps, time is in minutes, scale bar = 10µm.

**Video 6.** | **FIP localization to interzonal MTs in anaphase and telophase is independent of EB1.** Corresponds to Figure S4E. Cultured S2 cell transiently expressing mNG::FIP^Δ1-2^. Video begins in metaphase, where FIP^Δ1-2^ is concentrated as expected around mitotic chromatin, and extends through late telophase, where FIP concentrates as expected around the cell midbody. Image series is maximum intensity projection of six 0.8µm slices, acquired at 20 s/frame and displayed at 10 fps, time is in minutes:seconds relative to NEB, scale bar = 5µm.

**Video 7. | Low expression of Feo does not bundle MTs and does not interfere with FIP MT +end tracking in interphase.** Corresponds to Figure S6B. Interphase S2 cell coexpressing low quantities of mNeonGreen::Feo (red) and TagRFP::FIP (green) illustrating normal FIP MT +end tracking. Images are single planes acquired at 3 seconds/frame and displayed at 30 fps.

**Video 8. | FIP and Feo localization is indistinguishable during anaphase and telophase.** Corresponds to Figure 6C. Mitotic S2 cell coexpressing low quantities of mNeonGreen::Feo (red) and TagRFP::FIP (green) illustrating normal FIP MT +end tracking. Images are single planes acquired at 15 seconds/frame and displayed at 15 fps.

**Video 9. | Feo localization in wild-type peripodial wing disc cells.** Corresponds to Figure 7. Peripodial cells expressing Feo::GFP indicating the high level of mitotic activity. Feo localizes to interzonal MTs at anaphase and telophase, followed by the focused localization to the midbody. Images acquired at 20s/frame and displayed at 15 fps.

**Video 10. | FIP mutants have fewer mitotic divisions in wing disc cells.** Corresponds to Figure 7. Peripodial cells expressing Feo::GFP in *fip-* mutant animals highlighting the low level of mitotic activity. Localization of Feo to anaphase interzonal MTs is rarely observed, while Feo localization to midbodies is minimal. Images acquired at 20s/frame and displayed at 15 fps.

